# Controlling interfacial protein adsorption, desorption and aggregation in biomolecular condensates

**DOI:** 10.1101/2024.10.20.619145

**Authors:** Brent S. Visser, Merlijn H. I. van Haren, Wojciech P. Lipiński, Kirsten A. van Leijenhorst-Groener, Mireille M.A.E. Claessens, Marcos V. A. Queirós, Carlos H. I. Ramos, Jorine Eeftens, Evan Spruijt

## Abstract

The aggregation of amyloidogenic proteins is linked to age-related diseases. The presence of interfaces can affect their aggregation mechanism, often speeding up aggregation. α-Synuclein (αSyn) can adsorb to biomolecular condensates, leading to heterogenous nucleation and faster aggregation. Understanding the mechanism underlying localization of amyloidogenic proteins at condensate interfaces is crucial for developing strategies to prevent or reverse their binding. We show that αSyn localization to the surface of peptide-based heterotypic condensates is an adsorption process governed by the protein’s condensate-amphiphilic nature. Adsorption occurs in multiple layers and levels off at micromolar concentrations. Based on these findings, we design three strategies to modulate αSyn accumulation: (i) addition of biomolecules that decrease the condensate ζ-potential, such as NTPs and RNA, (ii) competitive adsorption of proteins targeting the condensate interface, such as G3BP1, DDX4-YFP, EGFP-NPM1, Hsp70, Hsc70, and (iii) preferential adsorption of αSyn to membranes. Removing αSyn from the condensate interface slows aggregation, highlighting potential cellular control over protein adsorption and implications for therapeutic strategies.

**Graphical abstract:** 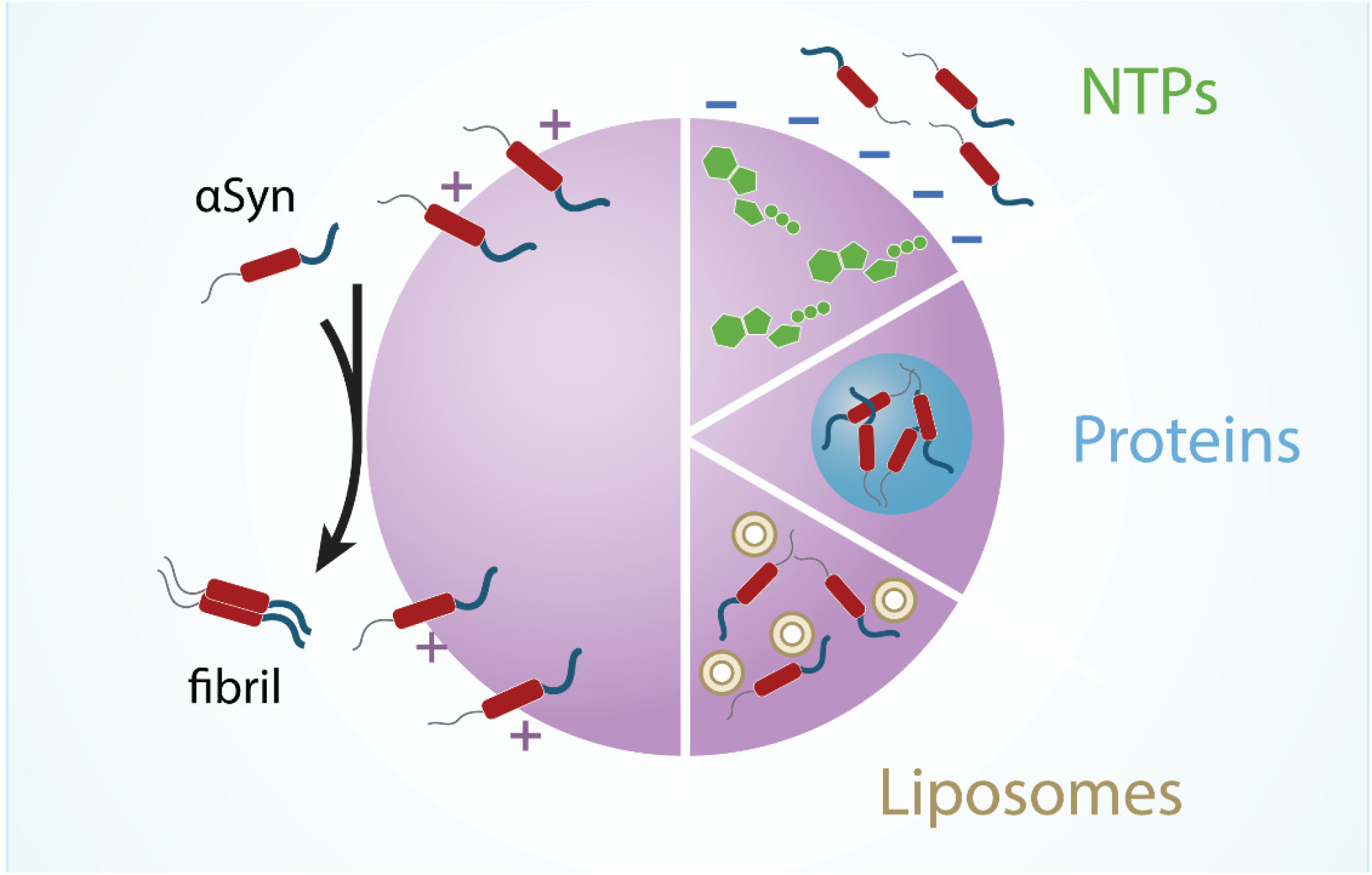

## Introduction

Cells constantly monitor the state of their proteome to ensure that misfolded or aggregated proteins are refolded or cleared. The accumulation of aggregated proteins can lead to severe cellular dysfunction, and ultimately result in diseases such as Alzheimer’s and Parkinson’s disease^1–5^. Interfaces can influence protein aggregation by serving as catalysts for heterogeneous nucleation. The influence and importance of interfaces on aggregation of proteins related to neurodegenerative diseases has been studied in the past^6^, using solid surfaces^7^, lipid membranes^8^, and water-air interfaces^9,10^. More recently, biomolecular condensate interfaces were found to also accumulate these proteins, and nucleate protein aggregation at their interface^11–13^. Linsenmeier et al. showed that formation of hnRNPA1 fibrils on idemic condensates consisting of hnRNPA1 takes place at the condensate interface, and that this process could be reduced by introducing protein-based surfactants^13^. Shen at al. observed that liquid-to-solid transition of FUS condensates is also initiated at their interface^12^. He et al. showed that FUS aggregates grow on the condensate’s surface^14^. However, the mechanism underlying interfacial localization remains poorly understood, and general strategies to prevent or reverse interfacial protein localization are lacking.

Biomolecular condensates have a non-zero surface charge, either because they are composed of charged biomolecules^15^, or because of asymmetric binding of ions^16,17^. Many disordered, aggregation-prone proteins have charged patches, such as α-synuclein (αSyn)^11^, tau^18^, FUS^19^, prion protein^20^, and TDP-43^21^, which could lead to attraction to the charged interface of the condensates. We hypothesize that surface charge of condensates can govern the interfacial localization of (disordered) proteins, and thus holds a key to preventing or reversing localization and heterogeneous nucleation-based aggregation.

Here, we study how model condensates with tunable ζ-potential accumulate wild-type αSyn at their interface, and how it can be controlled. We specifically investigate how the condensate ζ-potential is a driver in localization proteins, including αSyn and TDP-43, by measuring the charge of the interface prior to and after addition of the proteins. We show that protein adsorption follows a Freundlich-type adsorption isotherm, suggesting that the condensate interface exhibits heterogeneous binding sites arranged in multiple layers with a finite overall binding capacity. More importantly, it indicates that protein adsorption at condensate interfaces is an equilibrium process that can be reversed. Using these insights, we present three biologically relevant strategies to control αSyn interfacial localization and aggregation: (i) addition of biomolecules that can alter the ζ-potential of condensates, such as NTPs or RNA, (ii) competitive adsorption of proteins targeting the condensate interface, such as G3BP1, DDX4-YFP, EGFP-NPM1, Hsp70, and Hsc70, (iii) preferential adsorption of αSyn to membranes, sequestering them away from the condensate interface. These findings indicate that condensate ζ-potential and electrostatic interactions can govern accumulation of proteins at condensate interfaces and pave the way for strategies to control protein localization to condensate interfaces and prevent protein aggregation.

## Results and Discussion

### αSyn localization is governed by ζ-potential and protein amphiphilicity

We use model condensates with tunable ζ-potential, consisting of poly-D,L-lysine with 100 residues and poly-D,L-glutamate with 100 residues (further termed pLys/pGlu) to investigate the role of ζ-potential in the localization of amyloidogenic proteins at condensate interfaces. We recently showed that wild-type αSyn can be accumulated at the interface of pLys/pGlu condensates, leading to substantially enhanced rates of αSyn aggregation^20^. αSyn is a protein consisting of an active aggregation core, flanked by a positively charged disordered N-terminal fragment and a negatively charged disordered C-terminal tail (Fig. 1a)^11^. To gain more insight in αSyn interfacial localization, we first measured the localization of various αSyn mutants. Removal of the C-terminal domain drastically changes αSyn localization: the wildtype protein (computed pI=4.67; charge at pH 7.4 is -9.7) accumulates at the surface of pLys/pGlu droplets, while a variant without the negatively charged C-terminal domain, αSyn(1-108) (computed pI=9.16; charge at pH 7.4 is +2.29)^22,23^, was excluded from both the surface and the droplet interior (Fig. 1b)^11^. We also studied the partitioning of αSyn(60-140), which lacks the disordered, positively charged N-terminal domain (computed pI=4.05; charge at pH 7.4 is -12.44). αSyn(60-140) partitioned into droplets but did not accumulate at the interface (Fig. 1b). αSyn is thus amphiphilic to condensates, containing charged patches that prefer the interior of the condensate (residues 1-59) and parts that prefer the dilute phase (residues 109-140), causing its interfacial accumulation^11^.

**Figure 1:**
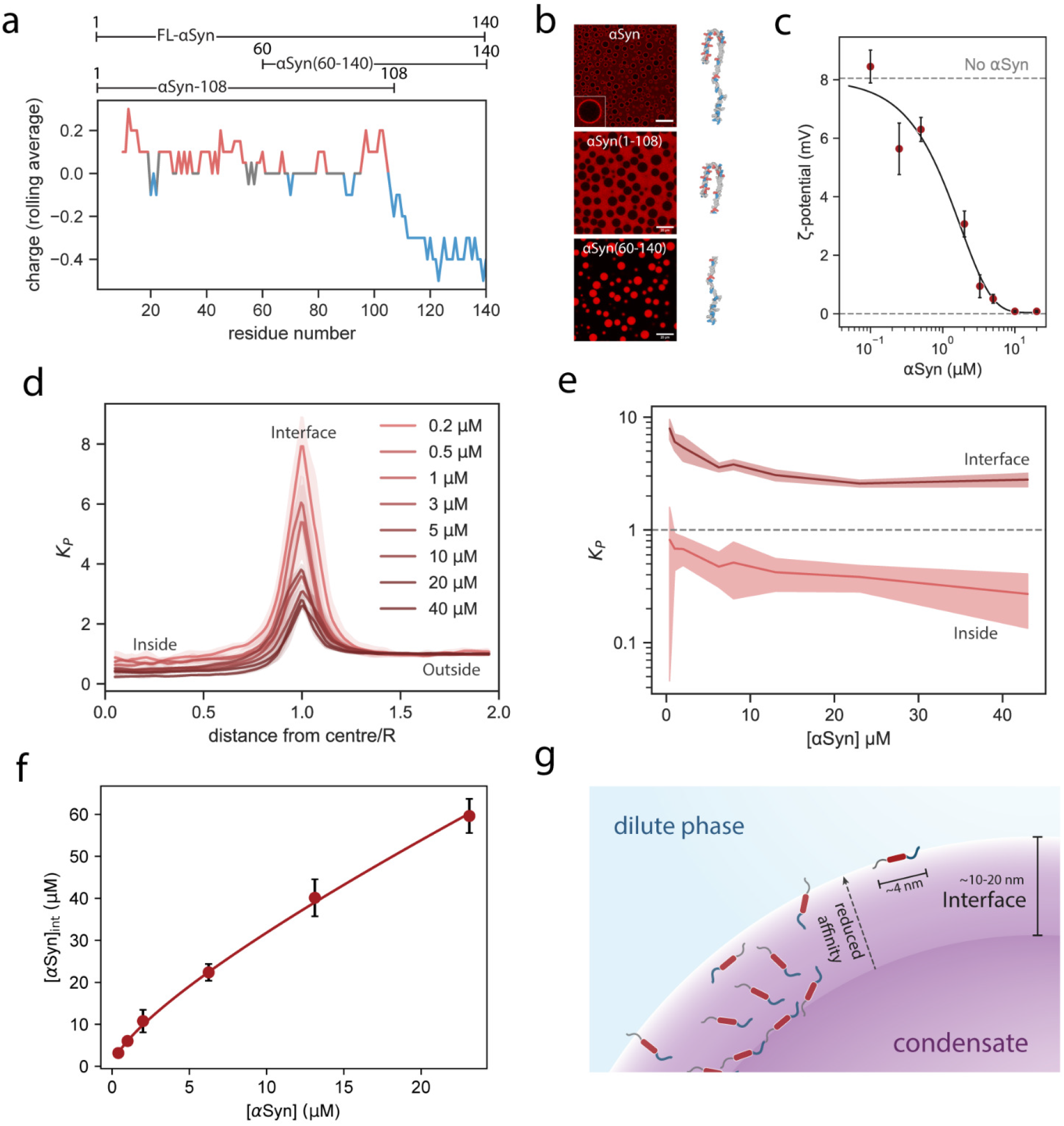
*α*Syn adsorbs to positively charged pLys/pGlu condensates and neutralizes the condensate *ζ*-potential. **(a)** Moving average of the charge distribution of αSyn and three variants used in this paper, αSyn, αSyn(60-140), and αSyn-108. 3D model of full-length αSyn bound to a micelle (PDB ID 1XQ8), negatively charged amino acids are shown in blue and positively charged amino acids in red^24^. The S9C variant was used for labeling. **(b)** αSyn is condensate-amphiphilic and depending on which domains of the protein are present can localize to the interface (WT), dilute (1-108), or condensed phase (60-140). **(c)** Addition of αSyn to pLys/pGlu condensates reduces the condensate ζ-potential in a concentration-dependent manner. Error bars indicate the standard deviation (*n* > 30 individual droplets). A sigmoidal fit of the data is shown. **(d)** Average αSyn partitioning profile of the condensate. Average of radial intensity profile of 6 droplets. **(e)** Partitioning of αSyn and interfacial binding for different concentrations of αSyn. Partitioning to both interface and inside is reduced upon addition of more αSyn. Shaded regions indicate the standard deviation. **(f)** Microscopically measured interfacial concentration of αSyn with Freundlich isotherm fitted to the datapoints (*S*_sat_ = 83.7 ± 36.5 μM, *n* = 1.58 ± 0.14). αSyn concentrations differ from (d) and (e). **(g)** Schematic drawing of the interfacial region of the condensates with multiple binding sites for αSyn with different affinities.

We then assessed how the accumulation of αSyn at the surface of condensates affects their ζ-potential by microelectrophoresis^15^. pLys/pGlu condensates had a ζ-potential of +8.1 ± 0.9 mV, which decreased to zero upon addition of αSyn in a sigmoidal manner, characteristic of an adsorption isotherm (Fig. 1c). To investigate the mode of interfacial adsorption, we measured the partitioning of αSyn at different concentrations, keeping the concentration of Alexa Fluor 647-labeled S9C αSyn (Alexa-647-αSyn) constant when the total concentration was above 3 μM (Supplementary Fig. 2). From this, a partition coefficient (*K*_P_) was calculated, which shows that the relative concentration of αSyn at the interface decreases as the total concentration increases (Fig. 1d, e).

**Figure 2:**
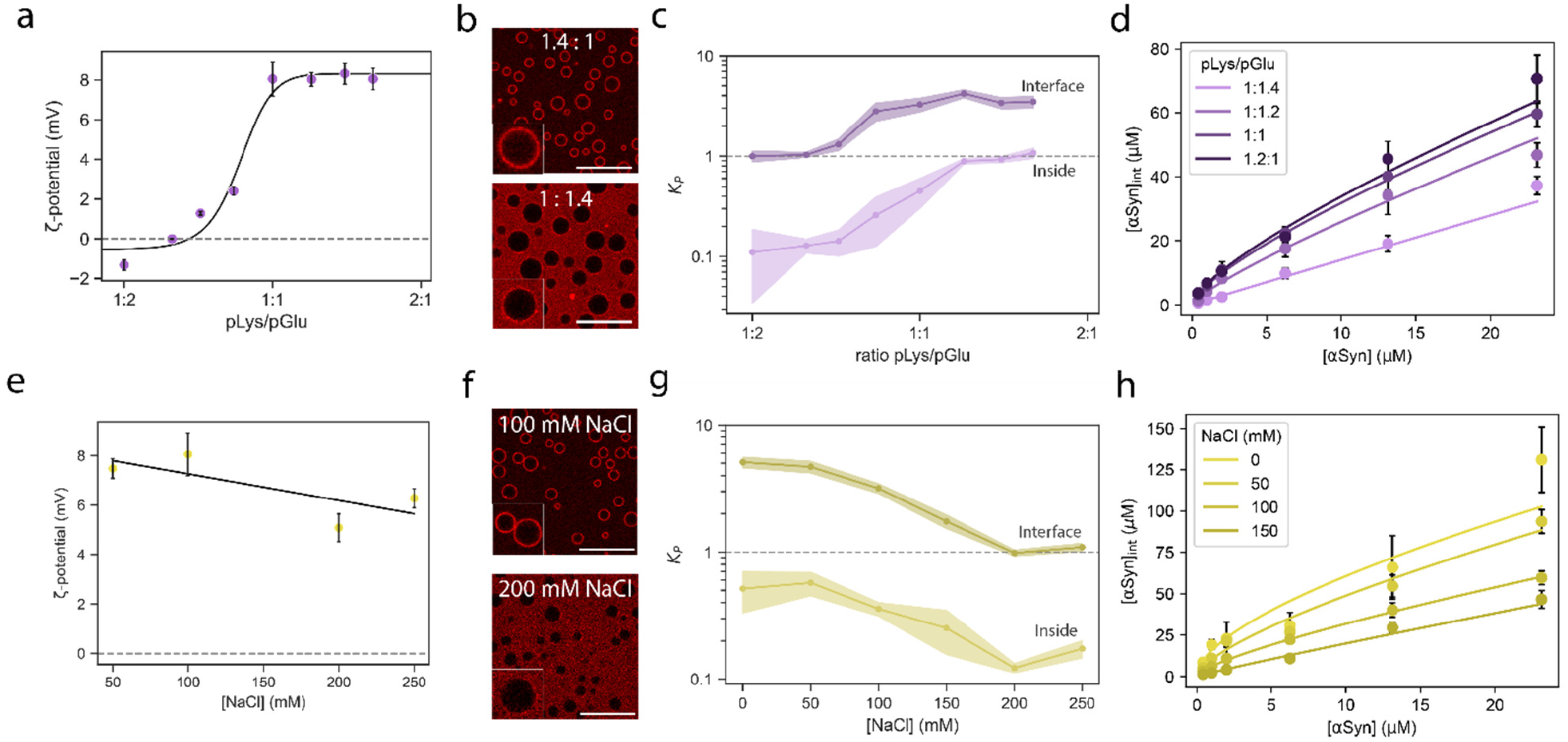
Changing the condensate *ζ*-potential and screening electrostatic interactions reduces the localization of *α*Syn on the coacervate interfaces. **(a)** The ζ-potential of pLys/pGlu condensates is dependent on the mixing ratio of polymers. Increasing the amount of pGlu or pLys decreases or increases the charge, respectively. Condensates are positively charged at neutral mixing ratio (1:1). **(b)** Confocal microscopy shows that αSyn accumulates at the interface of positively charged condensates but not at the interface of negatively charged ones. (Scale bar = 20 μm) **(c)** Partition coefficient of αSyn to the inside and surface of condensates for a range of pLys:pGlu ratios. **(d)** Microscopically measured interfacial concentration of αSyn with Freundlich isotherm fitted to the datapoints for 4 ratios of pLys/pGlu. **(e)** The ζ-potential of pLys/pGlu droplets slightly decreases with an increase in NaCl concentration. A linear fit of the data is shown. **(f)** Microscopy shows that an increase of NaCl decreases the interfacial accumulation of αSyn on the condensate interface. (Scale bar = 20 μm). **(g)** Partitioning of αSyn at the interface and to the interior of pLys/pGlu decreases with addition of NaCl. **(h)** Microscopically measured interfacial concentration of αSyn with Freundlich isotherm fitted to the datapoints for 4 concentrations of NaCl.

With increasing total concentration of αSyn, the amount of αSyn at the interface starts to level off, as can be seen upon plotting the interfacial concentration against total concentration of αSyn. We found that the amount of αSyn at the condensate interface is described best with the adapted Freundlich isotherm (Fig. 1f). Alternative adsorption isotherms, such as Langmuir or BET models performed worse as judged by the Bayesian information criterion (Supplementary Fig. 3 & 4). This suggests that the binding sites at the condensate interface are heterogenous and multilayered. We interpret this as a reflection of the presence of a transition region in between the condensate interior and the surrounding solution. Based on the magnitude of the condensate interfacial tension, the width of the condensate interfacial region of condensates is estimated to be large compared to the size of a single protein.

**Figure 3:**
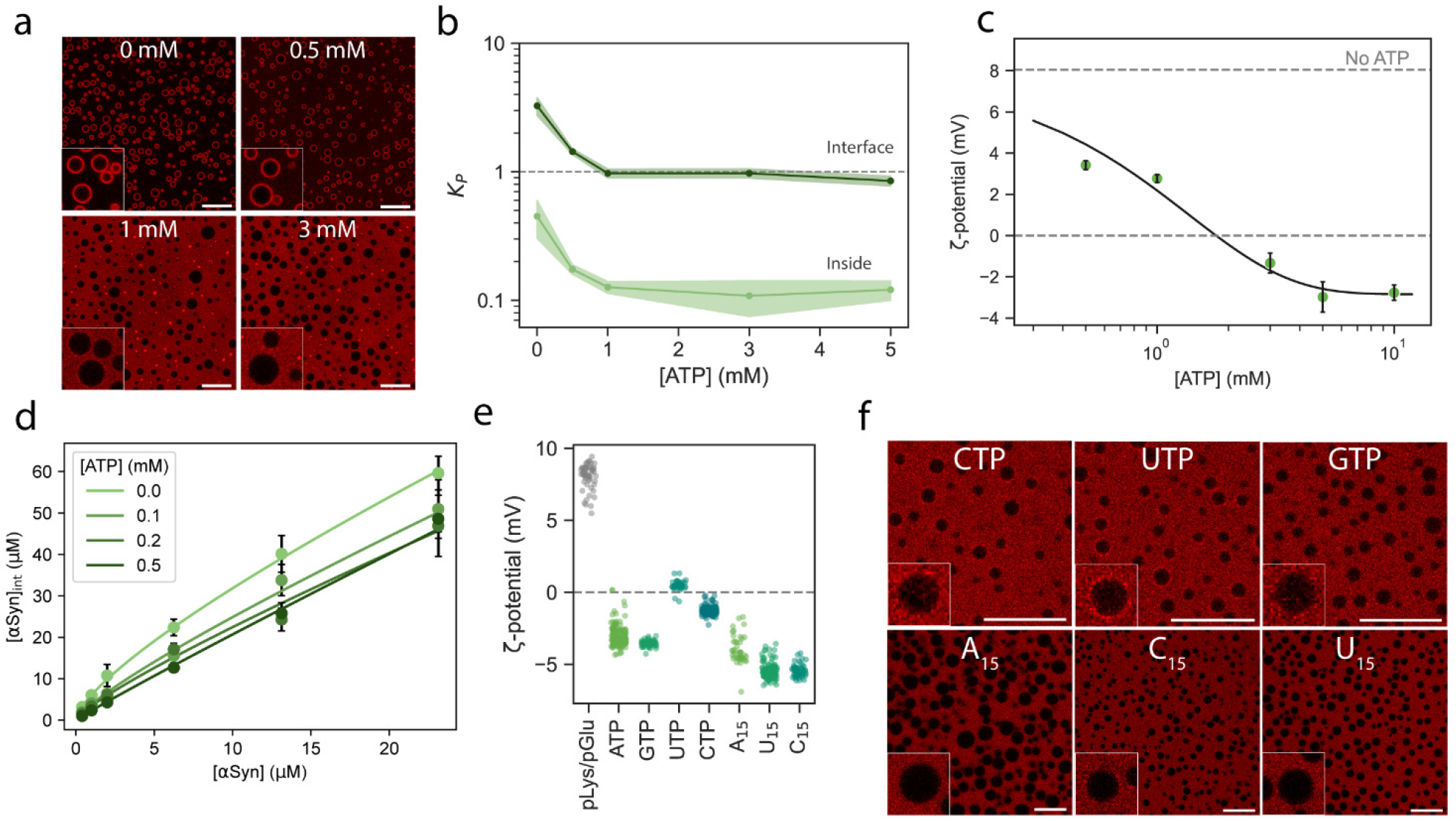
NTPs and RNA change the *ζ*-potential of pLys/pGlu condensates and decrease interfacial *α*Syn accumulation. **(a)** αSyn is delocalized from the pLys/pGlu condensate interface by addition of ATP. (Scale bar = 20 μm) **(b)** The *K*_P_ of αSyn is reduced by addition of ATP, both on the interface and inside of the condensates. **(c)** ATP causes a decrease in the ζ-potential of pLys/pGlu droplets. A sigmoidal fit of the data is shown. **(d)** Freundlich model fits of αSyn adsorption for different ATP concentrations. A decrease in the binding capacity of the surface can be seen as more ATP is added. **(e)** Addition of NTPs (5 mM) and RNAs (0.1 mM) reduce the ζ-potential of pLys/pGlu condensates. **(f)** NTPs (5 mM) and RNA oligos (0.1 mM) lead to removal of most of αSyn from the interface. (Scale bar = 20 μm)

**Figure 4:**
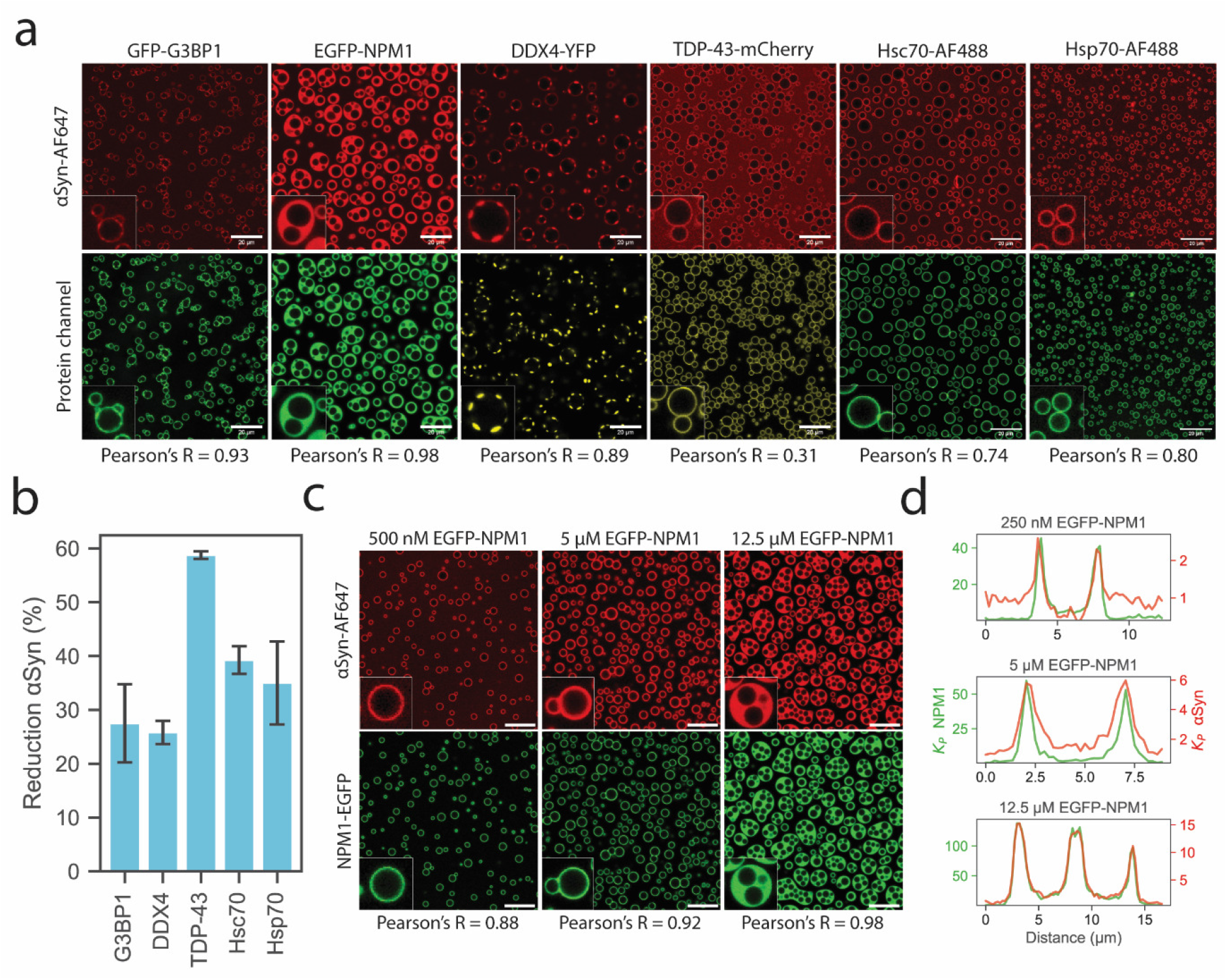
Localization of various proteins to the interface of pLys/pGlu condensates with *α*Syn. **(a)** GFP-G3BP1, EGFP-NPM1, DDX4-YFP, and TDP-43-TEV-mCherry localize to the interface of the condensates without moving αSyn to the dilute phase. Notably, NPM1 and DDX4 form a distinct third phase around the condensates, which also sequesters αSyn. Pearson’s correlation coefficients are shown to quantify colocalization. **(b)** Percentage reduction of the αSyn intensity at the interface in the presence of selected competing interfacial proteins. All proteins reduced αSyn intensity at the interface. (**c)** EGFP-NPM1 forms multiphase condensates at 12.5 μM, which sequester αSyn. (**d)** Line profiles of EGFP-NPM1 (green) and αSyn (red). The partition coefficient of both EGFP-NPM1 and αSyn increases with higher concentrations, and both colocalize in the multiphase condensates.

Across this interfacial region, the density and conformation of condensate components changes gradually^25,26^, and proteins such as αSyn can adsorb both close to the surrounding dilute phase where the interaction density is low, and deeper into the interfacial region, closer to the condensate bulk where the interaction density density is higher and the number of unbound sites that are available for binding may be lower (Fig. 1g). Taking the interfacial heterogeneity into account, the condensate interface has a finite overall capacity for adsorption, and the Freundlich isotherm allows us to compare the binding capacity of the condensates (reflected by the parameter *S*_sat_), as well as the characteristic interaction strength between the studied systems (reflected by *n*; see also Supplementary Information, Explanation of fitting of adapted Freundlich model).

Neutralization of the positive ζ-potential by negatively charged αSyn suggests that the adsorption is driven by electrostatic attraction. To test this hypothesis, we altered the condensate ζ-potential by changing the mixing ratio of pLys and pGlu. The ζ-potential of pLys/pGlu condensates at a 1:2 mixing ratio is negative (-1.3 ± 0.3 mV), at a 1.6:1 mixing ratio it is highly positive (+8.1 ± 0.5 mV, Fig. 2a). Theoretical work by Majee et al. showed that at equal molar ratios, charged condensate interfaces can occur due to unequal gradients of charged molecules at the interface, leading to a net charge, explaining the positive ζ-potential of pLys/pGlu condensates at 1:1 ratio^17^. We found that αSyn remained in the dilute phase for ratios lower than 1:1.4 pLys/pGlu (Fig. 2b, Supplementary Fig. 5). The relative amount of αSyn at the condensate interface and its interior, reflected by *K*_P_, increased with increasing ζ-potential and reached a plateau at the mixing ratio where the ζ-potential also reached a plateau (Fig. 2c). Fitting Freundlich isotherms to the data at different pLys/pGlu ratios shows that changing the ratio of condensate components alters the affinity of the surface to αSyn (Fig. 2d). Both the interaction strength and binding capacity increase drastically with increasing pLys (Supplementary Fig. 6).

**Figure 5:**
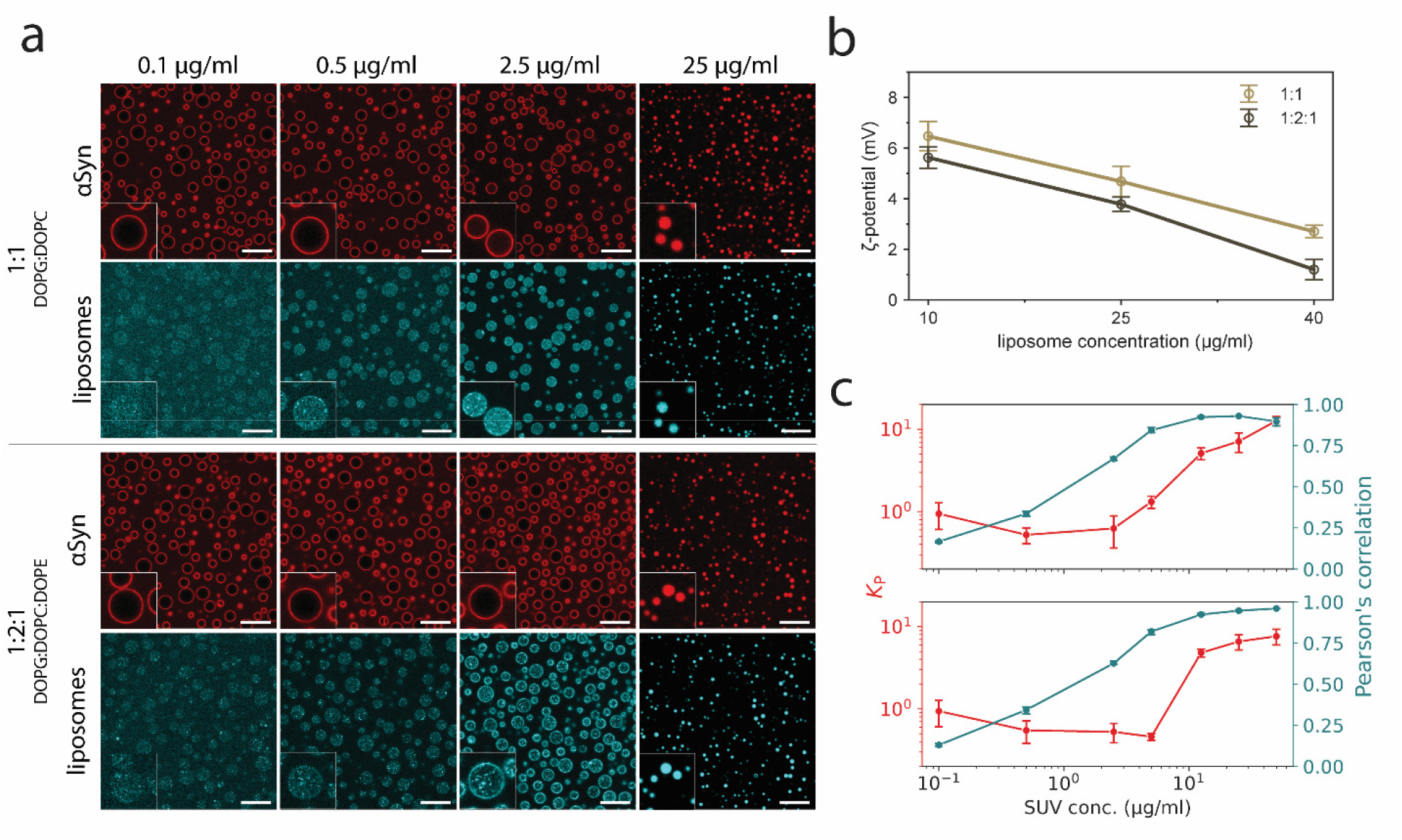
Liposomes partition into pLys/pGlu condensates and drive partitioning of αSyn. **(a)** Liposomes localize to the interface at low concentration and partition into the condensates at higher concentrations. This leads to co-partitioning of αSyn above a critical concentration. **(b)** ζ-potential of the condensates stays positive after addition of the liposomes. **(c)** Increasing liposome concentration leads to increased partitioning of αSyn (red) and colocalization of αSyn and the liposomes (blue).

**Figure 6:**
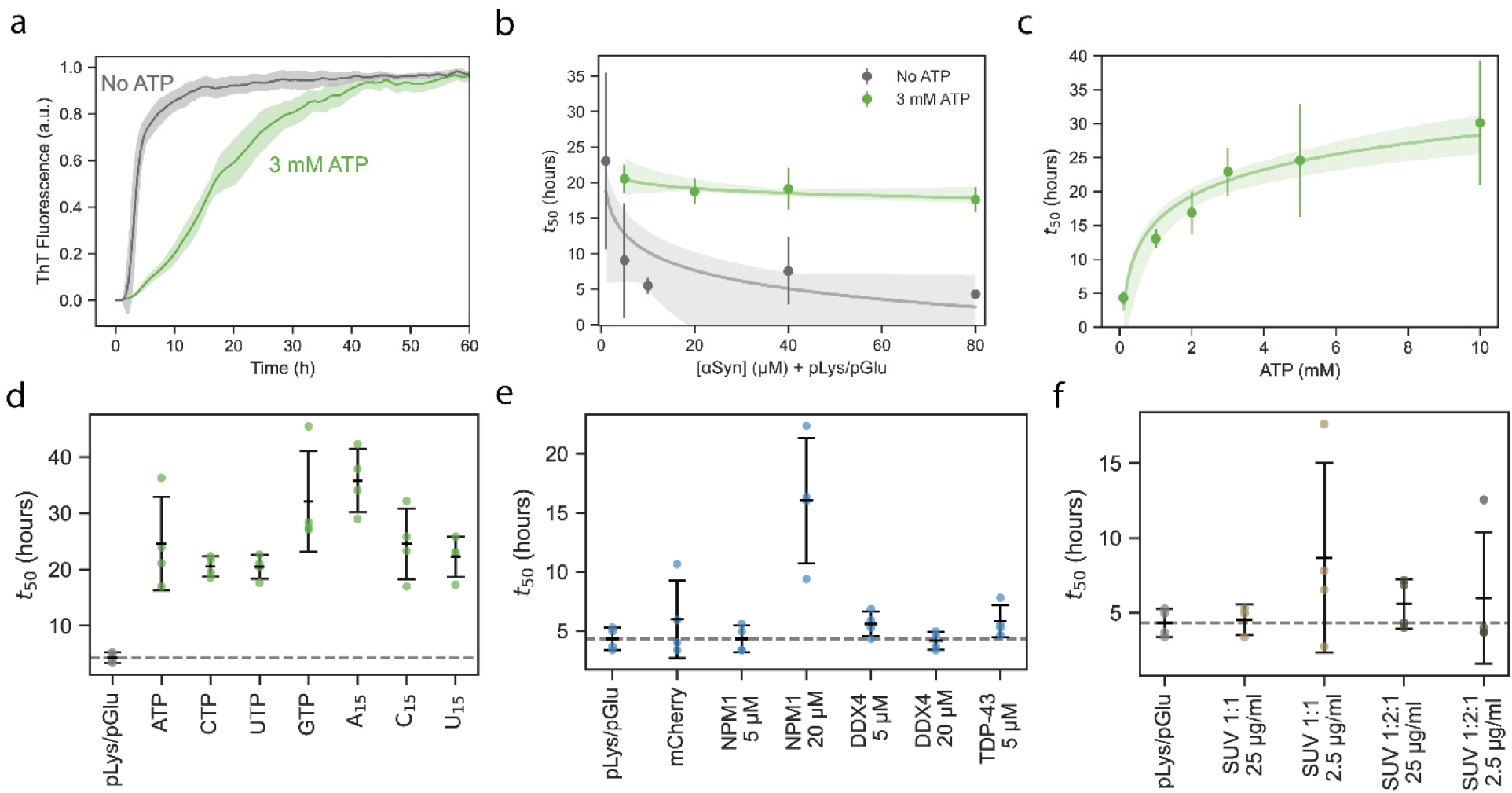
Removal of *α*Syn from the condensate interface with ATP slows down aggregation. **(a)** ThT aggregation assays show that the addition of 3 mM ATP to pLys/pGlu condensates substantially slows down aggregation (*n* = 4). **(b)** Increasing the concentration of αSyn in presence of pLys/pGlu condensates speeds up protein aggregation when no ATP is present but has no effect when 3 mM ATP is present. Exponential decay fits of the data are shown. **(c)** There is a logarithmic increase in the t_50_ of aggregation and concentration of ATP. **(d)** Aggregation of αSyn with pLys/pGlu condensates is slowed down by the presence NTPs (5 mM) or RNA oligos (0.125mM) compared to the reference sample. **(e)** Proteins that co-localize to the interface of pLys/pGlu condensates with αSyn do not substantially alter the aggregation rate. However, the multiphase condensates formed by NPM1 (20 μM) did slow down aggregation. **(f)** αSyn aggregation with pLys/pGlu condensates in presence of DOPG/DOPC 1:1 and DOPG/DOPC/DOPE 1:2:1 liposomes. The liposomes did not alter aggregation substantially for either concentration.

If αSyn localization is driven by electrostatic interactions, charge screening would weaken the attraction and reduce αSyn accumulation at the interface. Indeed, increasing NaCl only slightly reduced ζ-potential, but it drastically reduced αSyn adsorption at the interface (Fig. 2e-f). Above 250 mM NaCl, αSyn did not adsorb at all, while the *K*_P_ of αSyn inside the condensates decreased to approximately 0.15 (Fig. 2g). Freundlich fits show that increasing NaCl concentration also reduces both the binding capacity (Fig. 2h, *S*_sat_ = 333 μM to *S*_sat_ = 9.69 μM) and interaction strength (*n* = 1.86 to *n* = 1.15, Supplementary Fig. 6). We confirmed that this NaCl concentration could effectively screen the electrostatic interactions between αSyn and pLys by measuring the diffusion of fluorescently labelled αSyn bound to pLys with increasing NaCl concentration using Raster Image Correlation Spectroscopy (Supplementary Fig. 8)^27^. At 200 mM NaCl, αSyn diffusion in the presence of pLys was comparable to its diffusion without pLys, indicating that αSyn is no longer bound to pLys. We hypothesized that similar driving forces can also cause interfacial localization of other condensate-amphiphilic proteins with charged patches. We therefore also examined the partitioning of TDP-43-TEV-mCherry, which is known to exhibit disordered domains with a net negative and positive charge^28^, in combination with pLys/pGlu condensates. Similar to αSyn, TDP-43-TEV-mCherry preferentially partitioned to the interface of the droplets (Supplementary Fig. 9).

### NTPs and oligonucleotides can regulate condensate ζ-potential and remove αSyn

Our results indicate that condensate interfaces exhibit heterogeneous, multilayered adsorption sites, and that adsorption to these interfaces is an equilibrium process driven by charge complexation that can be reversed. Therefore, we set out to design strategies to regulate the adsorption and investigate the possibility of selective removal of αSyn from the interface. In a biological context, cells could (i) modulate condensate ζ-potential by regulating the levels of NTPs and metabolites, (ii) express proteins that compete for adsorption to condensate interfaces or form a new condensed phase at interface, or (iii) exploit the sequestration of proteins by other cellular interfaces, such as membranes.

We first tested the effect of NTPs, which are present at relatively high concentrations within cells, with ATP being the most abundant at concentrations of ∼2mM, surpassing the Michaelis constant of enzymes by two orders of magnitude^29–34^.We added ATP to pLys/pGlu condensates at physiologically plausible concentrations, and found that above 0.5 mM ATP, αSyn was excluded from the condensate interface, resulting in a three-fold decrease in *K*_P_ at the interface (Fig. 3a-b). Addition of ATP also caused a decrease in αSyn partitioning inside the condensate, with a *K*_P_ of 0.5 in the absence of ATP and 0.1 at ATP concentrations of 1 mM and higher (Fig 3b). The condensate ζ-potential also decreased from +8.1 ± 0.9 mV at 0 mM ATP to -3.0 ± 0.8 mV at 5 mM (Fig. 3c). This lower and negative ζ-potential weakens the electrostatic interactions of αSyn with the condensate surface, causing αSyn to be released into the surrounding dilute phase. Furthermore, αSyn localization is reversible, as most of the αSyn releases from the interface after adding ATP (Supplementary Fig. 10). We also observed that partitioning of αSyn(60-140) was not substantially affected by the addition of ATP, confirming that ATP mainly affects interfacial accumulation (Supplementary Fig. 1). Addition of ATP increased both the surface tension and viscosity (Supplementary Figs. 7, 11). Interfacial αSyn with increasing ATP can also be described by the Freundlich isotherm, reaching a *S*_sat_ value of ∼20 μM and *n* of 1.1 (Fig. 3d, Supplementary Fig. 6).

Besides ATP, we added other NTPs to see if they have a similar effect on the ζ-potential of pLys/pGlu condensates. Addition of 5 mM GTP led to droplets with a ζ-potential of -3.5 ± 0.2 mV, similar to ATP (Fig. 3e). The pyrimidines CTP and UTP had a weaker effect than the purines, but still lowered the ζ-potential. All NTPs prevented αSyn interfacial accumulation (Fig. 3f). Next, we set out to study the influence of short RNA oligonucleotides on the condensate ζ-potential, as their multivalency could enhance the charge reversal observed with NTPs. We selected A_15_, C_15_ and U_15_ and added them to pLys/pGlu condensates. As expected, oligonucleotides could also invert the ζ-potential of condensates at a concentration of 0.1 mM, and they were able to displace αSyn from the droplet interface (Fig. 3e-f).

We also compared the interfacial accumulation of TDP-43-TEV-mCherry on pLys/pGlu condensates with and without 5 mM ATP. Supporting our previous findings, ATP also decreased the accumulation of TDP-43-TEV-mCherry, although it did not completely remove the protein (Supplementary Fig. 9), possibly because TDP-43 was already partially aggregated.

The effect of low concentrations of NTPs and oligonucleotides on protein adsorption to condensates shows the potential to control protein localization based on electrostatic interactions, which could prevent heterogeneous nucleation at the condensate interface. These results suggest that electrostatic interactions may be a common factor in determining protein localization and that cells may use strategies like ATP production to manipulate condensate interface properties and protein localization.

### Proteins can displace αSyn by competitive adsorption

We then investigated if competitive adsorption at the condensate interface is possible, and if other proteins could displace αSyn from the interface. We investigated the condensate surface targeting potential of a number of commonly studied, partially disordered proteins with negatively charged patches (Supplementary Fig. 12), namely GFP-G3BP1, DDX4-YFP, EGFP-NPM1, and TDP-43-TEV-mCherry, and two heat shock proteins, Hsc70 and Hsp70, which are known to interact with αSyn^35–39^. Interestingly, most of these proteins were found to localize at the interface of pLys/pGlu condensates (Fig. 4a). The degree of displacement of αSyn (as measured by reduction in the *K*_P_ to the interface) varied between proteins, (Fig. 4b) and many co-localized with αSyn (Supplementary Fig. 13a), suggesting that their binding strength to the interface was comparable to αSyn. The molecular chaperones, Hsc70 and Hsp70, also co-localize with αSyn at the interface, potentially allowing for enhanced chaperone functionality^40–42^ and reducing the aggregation propensity of αSyn, without necessarily displacing it completely from the condensate interface. We also observe that the chaperones localize to the interface more strongly when αSyn is present, showing potentially targeted accumulation (Supplementary Fig. 14).

Surprisingly, addition of DDX4-YFP, GFP-G3BP1, and EGFP-NPM1 led to the formation of multiphase condensates when added to the pLys/pGlu condensates, resulting in αSyn partitioning in the newly formed outer phase with high colocalization. A reason for this additional phase separation is the amphiphilic nature of these proteins, as they contain a phase separating domain and a solubilizing domain in the form of the fluorescent tag^43^. For EGFP-NPM1, we observe multiphase condensate formation at 12.5 μM (Fig. 4c, Supplementary Fig. 13a), and high partitioning of αSyn into the new phase (Fig. 4d, *K*_P_ = 12). The colocalization of NPM1 and αSyn increases with NPM1 concentration (Supplementary Fig. 13b and 15). The multiphase condensates can potentially function as a compartment capable of sequestering αSyn to remove it from the condensate interface^11^. We found that the ζ-potential decreased with increasing EGFP-NPM1 and became negative around 2.5 μM, even before the separate new phase is visible by microscopy (Supplementary Fig. 15). This is remarkable, as αSyn was not able to invert the droplet ζ-potential. We hypothesize that EGFP-NPM1 already forms a new microphase that is spread on the interface of the pLys/pGlu condensates and that sequesters αSyn. This newly formed liquid phase has a negative ζ-potential, providing a separate mechanism of removing αSyn from the pLys/pGlu condensate interface.

### Membranes can sequester αSyn away from condensate interfaces

Lastly, we investigated αSyn localization in the presence of a second interface to which the protein could adsorb. Preferential adsorption to the second interface could lead to removal of αSyn from the condensate interface. The role of αSyn is thought to involve membrane binding^44^. Membrane-bound αSyn displays distinct conformations from monomeric αSyn^45^, sometimes leading to delayed aggregation and nonparticipation in aggregation^46^. In vitro, it has been shown that αSyn can also bind cooperatively to unilamellar liposomes^47,48^. We prepared vesicles with two lipid compositions, DOPG/DOPC 1:1 and DOPG/DOPC/DOPE 1:2:1 (Supplementary Fig. 16) and added them to the condensates with αSyn.

Using labeled liposomes, we observed that liposomes of both compositions partition to the interface at low concentrations, and into the condensates at higher concentrations (Fig. 5a). We found that the liposomes lowered the ζ-potential of the condensates, but that they remained net positively charged (Fig. 5b). However, αSyn no longer localized to the condensate interface when the liposomes partition into the condensates. In contrast to other experiments with such positive condensates, αSyn partitions into the condensates and colocalizes with the liposomes (Fig. 5c). We hypothesize that αSyn is sequestered by the membrane surface and is pulled into the condensates by the vesicles. It remains to be investigated whether the vesicles remain intact upon partitioning into the condensates. No substantial differences between the two compositions of liposomes was observed in the partitioning, despite a difference in ζ-potential of the droplets. These results show that competition between interfaces can alter protein localization, which could also serve as a regulatory mechanism in cells, as these interactions are likely protein and condensate specific.

### Displacement of αSyn from the condensate interface slows down aggregation

We have previously reported that condensate interfaces can act as heterogeneous nucleation sites for αSyn aggregation^11^. It has also been reported that positively charged interfaces accelerate aggregation^49^. We therefore expected that removal of αSyn from pLys/pGlu interface can affect the aggregation kinetics, and lead to a reduction in aggregation rates. We used Thioflavin T (ThT) fluorescence assays to follow αSyn aggregation with condensates in the different scenarios investigated above to remove αSyn from the condensate interface. Addition of 3 mM ATP substantially slows down αSyn aggregation in the presence of condensates (Fig. 6a). This effect is observed over a range of αSyn concentrations, from 5 to 80 μM (Fig. 6b). Increasing the ATP concentration could further slow down αSyn aggregation up to fivefold (Fig. 6c). While addition of ATP changes the aggregation half-time substantially in the presence of condensates, we find no major influence on the half-time in the absence of condensates, confirming that ATP acts as a modifier of αSyn-condensate interactions, rather than interacting with αSyn directly (Supplementary Fig. 17). As negative control, we examined aggregation of αSyn(60-140) – which does partition to the interface – and observe that aggregation is not substantially altered due to addition of ATP (Supplementary Fig. 18). These findings correspond to the observations of ATP concentration-dependent αSyn removal from the condensate interface (Fig. 3a).

Since we observed removal of αSyn not only for ATP but also for other NTPs and small RNA oligos, we investigated whether they could also reduce the aggregation rates. As expected, all of the strategies that removed αSyn from the interface also slowed down aggregation (Fig. 6d). One of the proteins was also able to slow down aggregation – 20 μM EGFP-NPM1 – by forming multiphase condensates that sequester αSyn (Fig. 6e). In the case of smaller multiphase compartments, we hypothesize that the physicochemical environment did not induce aggregation-slowing conformational changes in αSyn. We also measured the aggregation kinetics in the presence of liposomes. On average, the addition of liposomes had no substantial effect on the aggregation rate of αSyn. This can be explained by the fact that αSyn aggregation can also be enhanced by binding to membranes, similar to its binding to condensate interfaces (Fig. 6f).

We also examined the kinetics of αSyn aggregation with increased NaCl and altered condensate composition. Interestingly, while NaCl accelerates αSyn aggregation in solution^50^, it suppresses aggregation of αSyn in the presence of condensates (Supplementary Fig. 19), by removing αSyn from the interface^50^. Altering the condensate composition also influences αSyn aggregation rates. As expected, neutral or negatively charged condensates slow down aggregation, whereas positively charged condensates accelerate it (Supplementary Fig. 19).

Finally, we analyzed how nucleation rates and elongation rates change when αSyn no longer accumulated at the interface of the condensates. We calculated the lag time (*t*_lag_) and 1/*V*_max_ as kinetic parameters that can serve as proxies for *k*_n_ and *k*_elong_ using methods described previously^11^. We found that the aggregation lag time *t*_lag_ is increased in all situations when αSyn is removed from the interface, indicating suppression of nucleation (Supplementary Fig. 19). Interestingly, the maximum rate of aggregation is reduced by addition of NaCl and ATP, but not by altering the pLys/pGlu ratio.

## Conclusion

Increasing evidence shows the interface of biomolecular condensates influences protein aggregation. This applies to aggregating proteins that undergo LLPS within cells on their own, but also to systems in which the aggregating protein is a guest molecule in host liquid condensates formed by other components of the cell. To assess their relevance in pathology and to design strategies to prevent or reverse protein binding to condensate interfaces, we must first understand how the complex intracellular environment affects the properties of liquid interfaces and modulates interactions with aggregating proteins.

In this work we examined the interactions between a tunable model peptide-based condensate system, pLys/pGlu, and the amyloidogenic protein αSyn. αSyn binds strongly to the condensate interface when it carries a net positive charge, as observed with microscopy and ζ-potential measurements. The interface accumulation is an equilibrium adsorption process that follows a Freundlich isotherm, indicating a multilayered interface with heterogeneous binding sites. We confirmed that the interfacial charge of pLys/pGlu condensates and the amphiphilic nature of αSyn govern the adsorption.

We then proposed three biochemically relevant strategies to modulate or reverse αSyn adsorption to condensate interfaces and studied their effect on the kinetics of αSyn aggregation. NTPs and short oligonucleotides decreased the interfacial charge even at low concentrations, and could remove αSyn from the condensate interface in a reversible manner. Adding competitively adsorbing proteins, such as EGFP-NPM1, DDX4-YFP and GFP-G3BP1, was found to result in a reduction of αSyn at the interface and the formation of multiphase condensates, providing a more favorable phase for αSyn to partition into. Molecular chaperones Hsc70 and Hsp70 co-localize with αSyn at the interface of pLys/pGlu condensates, without necessarily displacing αSyn completely from the condensate interface. Finally, liposomes provided an alternative surface for αSyn to which it had a higher affinity, resulting in sequestration of αSyn away from the interface inside the condensates.

For some of these strategies, we observed substantially reduced αSyn aggregation rates. Adding NTPs and oligos slowed down aggregation up to fivefold, while NTPs did not affect αSyn in solution. In addition, EGFP-NPM1 multiphase condensates also slowed down αSyn aggregation. In contrast, addition of liposomes did not change the rate of αSyn aggregation, possibly because aggregation can also be enhanced by membrane binding. In summary, our study enhances understanding of protein interactions with the surface of condensates, revealing mechanisms cells might use to control aggregation against heterogeneous nucleation, through regulation of metabolite levels, and expression of RNA and proteins.

## Materials and methods

### Condensate formation

All experiments with poly-D,L-lysine and poly-D,L-glutamate condensates were performed using a HEPES buffer (final concentration 50 mM, pH 7.4) containing 100 mM NaCl and 100 μM EDTA unless stated otherwise. Condensates were prepared by adding pLys to the buffer, followed by pGlu, both with a final concentration of 2.4 mM monomer units. Additives such as sodium chloride, αSyn and ATP were added directly after, and the samples were mixed by vortexing for 10 seconds at 2800 rpm (lab dancer, VWR).

### *α*Syn preparation and labeling

Wild-type αSyn, and the cysteine mutants were expressed and purified as previously described^51^. Purified proteins were stored at a concentration of ∼250 μM in 10 mM TRIS-HCl (pH 7.4) at −80 °C, supplemented with 1 mM dithiothreitol (DTT) for the cysteine mutants. Single labeled proteins were labeled according to the dye manufacturer procedures.

### Protein expression

pHBS834 H14-SUMO-TDP43 WT-TEV-mCherry was a gift from Rajat Rohatgi (Addgene plasmid # 133320 ; http://n2t.net/addgene:133320 ; RRID:Addgene_133320).^28^ E. coli BL21 (DE3) was transformed with pHBS834 H14-SUMO-TDP43 WT-TEV-mCherry. Overnight cultures were used to inoculate large flasks of LB media, then cells were grown at 37 °C to an OD_600_ of 0.5, before protein expression was induced with 50 µ M IPTG overnight at 15 °C. Prior to harvesting, the pellet was resuspended in lysis buffer (20 mM Tris-HCl pH 8.0, 1000 mM NaCl, 5 mM DTT, 20 mM imidazole) with a cOmplete™, EDTA-free Protease Inhibitor Cocktail.

The resuspended cells were lysed using a homogenizer. The cleared supernatant was loaded onto a 5 ml Cytiva HisTrapFF. The His-tagged protein was eluted using elution buffer (20 mM Tris-HCl pH 8.0, 1000 mM NaCl, 5 mM DTT, 400 mM imidazole). ULP protease was added to cleave the His-SUMO-tag and was dialyzed overnight in a 25 kDa membrane against SEC buffer (40 mM HEPES pH 7.4, 300 mM NaCl, and 1 mM DTT). Finally, the protein was isolated SEC using a S200 16/600 SEC column. The protein was concentrated to approximately 100 μM using 10 kDa spin filters.

Human Hsp70 and Hsc70 were expressed in *Escherichia coli* BL21 (DE3) and Rosetta^TM^ 2(DE3) cells, respectively. The bacteria were left to grow until optical density has reached 0.6 – 0.8. At this point, 1 mM of IPTG was added, inducing the bacterial culture for 16 h at 16 °C. Cell pellets were obtained by centrifugation and stored at –20 °C. To be lysed, cells were resuspended in appropriate volumes of a buffer containing 20 mM phosphate, 250 mM NaCl, and pH 7.5 (buffer A) + 6 units of DNAse and RNAse, and 1 capsule of cOmplete™ for 30 min. After that, the suspensions were submitted to ultrasound pulses for a total time of 2 min and followed by centrifugation. Ni-affinity chromatography was performed by loading the lysed solutions onto a 5 ml HisTrap^TM^ columns (Cytiva). The elution of the proteins took place by employing the buffer A contaning 250 mM imidazole. The fractions containing each chaperone were loaded onto a Superdex^TM^ 200 pg HiLoad^TM^ 16/600 (GE healthcare) also equilibrated with buffer A. After that, the fractions enriched in the chaperones were analyzed through SDS-PAGE and concentrated by using a centrifugation filter with 30 kDa cutoff.

EGFP-NPM1, a fusion protein of enhanced green fluorescent protein and nucleophosmin-1 was expressed using BL21(DE3) *E. coli* cells transformed with a pET28a(+)EGFP-NPM1 plasmid. Cells were grown at 37 °C to an OD_600_ of 0.7 expression was induced with 1 mM IPTG. Cells were lysed using three rounds of sonication (Sanyo Soniprep 150) on ice in cycles of 10 s with an amplitude of 10%. EGFP-NPM1 was purified using a Ni-NTA agarose (Fisher Sci) column, followed by size exclusion chromatography with a HiLoad Superdex 75 26/600 (GE healthcare)^52^.

GFP-G3BP1 was expressed using U2OS cells that express GFP-G3BP1 at endogenous G3BP1 levels. Cells were lysed using short mild sonication on ice. GFP-nanobodies attached to agarose beads were used to purify GFP-G3BP1 in a one-step purification. An acidic glycine buffer (pH 2.4) was used to elute GFP-G3BP1, the solution was neutralized using Tris base (pH 10.4)^52^.

DDX4-YFP, a mutated version of the human DDX4 nuage protein, in which its C-terminal helicase domain is replaced by a yellow-fluorescent protein sequence was expressed using BL21(DE3) *E. coli* cells transformed with a Ddx4N1YFP pETM30 plasmid. Cells were lysed using a homogenizer (LTD FPG12800) and clarified cell lysate was purified using a HisTrapFF (Cytiva) column. TEV protease was added to the eluted DDX4-YFP protein and the mixture was dialyzed to SEC buffer (20 mM Tris, 300 mM NaCl, 5 mM TCEP, pH 8.0). The dialyzed product was concentrated and further purified using size-exclusion chromatography with a S200 16/600 SEC column (GE healthcare)^53^.

### Lipids and liposome preparation

All unlabeled lipids, 1,2-dioleoyl-sn-glycero-3-phosphoglycerol (DOPG) 1,2-dioleoyl-sn-glycero-3-phospho-ethanolamine (DOPE), and 1,2-dioleoyl-sn-glycero-3-phosphocholine (DOPC), were purchased from Avanti Polar Lipids. Atto 488-labeled DOPE was obtained from Sigma Aldrich. Lipid stock solutions were prepared by dissolving in chloroform at 25 mg/ml, evaporating, and re-dissolving in half of the original volume of chloroform for a final concentration of 50 mg/ml for the lipid stock solutions^54^.

DOPG/DOPC 1:1 and DOPG/DOPC/DOPE 1:2:1 liposomes were prepared according to^59^. Lipid solutions were added to chloroform in glass HPLC vials. The mixes were evaporated under argon to dry to a film. Buffer (50 mM HEPES, 100 mM NaCl, 5 μM EDTA) was added to each vial. Vials were kept at 40°C overnight. After incubation, contents were vortexed and extruded 11 times with a 200 nm membrane and 11 times with a 50 nm membrane.

### Preparation of modified glass slides

Samples were imaged in µ-Slide 18 Well chambered coverslips (uncoated polymer coverslip, Ibidi). All slides used for microscopy were modified to minimize spreading of the condensates on the surface of the slide. The surface intended to be modified was cleaned with oxygen plasma, and a solution of 0.01 mg/ml PLL-g[3.5]-PEG (SuSoS, Dübendorf, Switzerland) dissolved in 10 mM HEPES buffer (pH 7.4) was applied on the glass immediately after the plasma treatment. Glass was incubated with the PLL-g-PEG solution overnight at room temperature. Subsequently, it was rinsed three times with Milli-Q water and dried with pressurized air. Modified slides were stored at room temperature.

### Confocal microscopy

Localization of labeled proteins was studied using confocal microscopy. A Leica SP8x confocal microscope equipped with ×100 magnification oil-immersion objective was used. Samples were placed in 18-well chambered glass coverslips (Ibidi GmbH, Germany), previously modified with PLL-g[3.5]-PEG. Partition coefficients were determined by calculating ratio of fluorescence intensity in the condensed phase to fluorescence intensity in the outer phase (average intensity values from at least 6 droplets and from outer phase of similar area were used). Fluorescence profiles were measured using the ImageJ plugin Radial Profiles Extended and normalized to dilute phase signal. Colocalization was measured using the ImageJ plugin Coloc 2.

### Raster Image Correlation Spectroscopy

The diffusion of αSyn was determined using Raster Image Correlation Spectroscopy (RICS) on a Leica SP8 confocal microscope equipped with a single-photon detector. Calibration of the focal volume waist ω_0_ was performed using the known diffusion coefficient of Alexa 488 of 435 μm^2^ s^−1^ (*T* = 22.5 ± 0.5 °C) in water^55^, and ω_z_ was set to 3 times the value of ω_0_. All measurements were captured at a resolution of 256 × 256 pixels with a 20 nm pixel size using a 63x objective. Analysis of autocorrelation curves was performed in PAM.^56^

### *ζ*-potential measurements by microelectrophoresis

All samples were imaged on 6-well µ-channel slides (Ibidi) that were modified with 0.1 mg/ml PLL-g[3.5]-PEG. Before image acquisition, a 100 µ L condensate suspension was transferred to the channel and was incubated for 1 h to allow droplets to coalesce and settle on the glass surface. Electrodes (2 mm, silver) connected with copper wires to a BT-305A PSU direct current power source (Basetech) were lowered into opposing ends of the microchannel slide and an electric field of 1.2 to 12 V/cm was applied, with the cathode at the top of the field of view. Moving condensates were imaged in the middle of the channel of the microslide. Samples were imaged on an Olympus IX83 inverted fluorescence microscope equipped with a motorized stage (TANGO, Märzhäuser) and LED light source (pE-4000 CoolLED). Images were recorded with a 40× universal plan fluorite objective (WD 0.51 mm, NA 0.75, Olympus) with a temperature-controlled CMOS camera (Hamamatsu Orca-Flash 4.0).

Raw microscopy videos were processed and analyzed with MATLAB 2021 Image processing Toolbox and droplet trajectories were determined using methods previously described^15^. ζ-potentials for all detected droplets in a sample were determined from their velocities with a modified Smoluchowski equation, using the applied electric field strength, Debye length calculated from salt concentration and the droplet viscosity determined by active rheology. All parameters used to calculate the condensate ζ-potential are available in Supplementary Table S1.

### ThT aggregation kinetic assays

To estimate the kinetic parameters of aggregation, we performed standard thioflavin T (ThT) aggregation assays. Upon binding to β-sheets, ThT fluorescence intensity and the changes in fluorescence are proportional to the mass of aggregate formed^57,58^.

Aggregation assays were performed under the following conditions unless mentioned otherwise: 50 mM HEPES (pH 7.4), 100 mM NaCl, 100 μM EDTA, 20 μM ThT, and 40 μM αSyn. Protein solutions were filtered using Pierce cellulose acetate filter spin cups (Thermo Fisher Scientific) before every aggregation kinetic assay, and concentration was determined on the basis of absorbance at 276 nm (ϵ = 5600 M^−1^ cm^−1^ for wild-type αSyn). All aggregation assays were performed in non-binding 384-well black-walled plates (Greiner Bio-One GmbH, Austria) at 37°C. To prevent evaporation, wells in the two outer rows were always filled with water and the plate was sealed with a transparent sticker. Measurements were performed using a Tecan Spark microplate reader. Fluorescence intensity was recorded every 6 minutes using the bottom readout with continuous linear shaking in between. The excitation and emission wavelength range were controlled using filters (430 nm with 20 nm bandwidth and 460 nm with 20 nm bandwidth, respectively). Four measurements were done for every individual data point. To extract *t*_50_ from the ThT fluorescence traces, we fitted a simple aggregation model (as described previously^11^ and used the time to reach 50% of the max signal.

### NMR spectroscopy

To determine the partition coefficient of ATP, a 10 ml condensate suspension containing 2.4 mM pLys, 2.4 mM pGlu and 0.5 mM ATP in standard buffer with 10% D_2_O was centrifuged for 30 minutes at 500 g at 20 °C. The phases were separated and 12.5 µL condensate phase was obtained, which was diluted 40 times to 500 µL using 1 M NaCl. The supernatant was used without further dilution. Subsequently, hexamethylphosphoramide (HMPA) was added to the separated phases, with a final concentration of 4 mM for the condensate phase sample and 10 mM for the dilute phase sample. ^31^P NMR spectra were recorded on a Bruker-Avance III 500 spectrometer at 500 MHz. A pulse sequence was set up with 8 transients (nt = 8), P1 = 13 ms, which corresponds to approximately a 90° pulse angle and a d_1_ relaxation delay of 30 s, in order to ensure full relaxation of nuclei. The data was processed with MestReNova 14. The ATP concentration was calculated by taking the mean integral of the ? (-5.5 ppm), β (-10.5 ppm) and γ (-20.7 ppm) phosphate peaks. An ATP concentration of 101.2 mM and 0.22 mM was calculated in the condensate phase and dilute phase, respectively, resulting in a partition coefficient of 462.

### Fusion of suspended droplets using optical traps

Fusion assays in optical traps were performed to determine the inverse capillary velocity of condensates, based on protocols from^60^ and^61^. Fusion events were tracked using a LUMICKS C-Trap dual-trap OT instrument. Measurements were performed in Ibidi single-channel slides (µ-Slide I Luer, 0.4 mm, polymer bottom). Channel slides were modified with PLL-g[3.5]-PEG using the same protocol as for the glass slides, but using 0.003 mg/ml PLL-g[3.5]-PEG concentration, which is important for the further active rheology measurements performed in the same slides. The condensate samples were first mixed in an Eppendorf tube, then the suspension was transferred into the channel slide and placed at the OT instrument. Both traps were first set to intermediate power with ca. 30 µm distance between them to scavenge nearby condensate droplets. When the trapped droplets reached the desired size (5-10 µm in diameter), the droplet in trap 1 was moved close to the droplet in trap 2. Subsequently, trap 1 was moved at constant speed of 0.1 µm/s in the direction of trap 2 until fusion of the droplets was observed. The force-time response from both traps was recorded at 78.125 kHz sampling frequency and analyzed using the following model:

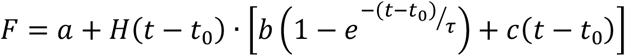

Where *a* and *a* + *b* are the pre-fusion and post-fusion force plateau, *t*_0_ is the starting time of the fusion event, *H*(*t* − *t*_0_) is a Heaviside step function applying the exponential term only after time *t*_0_, *τ* is the fusion relaxation time and *c*(*t* − *t*_0_) is a linear term compensating for the trap movement.

### Active rheology using an optical trap

Active rheology inside condensate droplets to measure interfacial tension and viscosity of condensates was performed by oscillating polystyrene beads, based on the protocol from^62^. After condensate droplets sedimented to the bottom of the channel slide and coalesced into larger droplets, 1 µl of Fluoresbrite Yellow Green Microspheres (1 µm diameter) suspension was added to the slide. Beads were either trapped in solution and dragged into the droplets, or beads that were already present inside droplets were used.

## Acknowledgements

This work was supported financially by a Vidi grant from the Netherlands Organization for Scientific Research (NWO), the European Research Council (ERC) under the European Union’s Horizon 2020 research and innovation program under grant agreement number 851963, and the São Paulo Research Foundation (FAPESP), under project number 2023/12135-0.

We would like to thank Dr. Alain A.M. André for purification of EGFP-NPM1 and GFP-G3BP1, Prof. Sushma N. Grellscheid for providing U2OS cells expressing GFP-G3BP1, Dr. N. Amy Yewdall for purification of DDX4-YFP, Jonathan Hoekstra for help in purifying TDP-43-TEV-mCherry, and Dr. Tim Nott for sharing the Ddx4N1YFP pETM30 vector.

## Author contributions

B.S.V, W.P.L, M.H.I.vH, and E.S. conceived the project. B.S.V., W.P.L., and M.H.I.vH. designed and performed the experiments. K.A.vL, MM.A.E.C, M.V.A.Q, J.E., and C.H.I.R, provided resources. B.S.V, W.P.L, M.H.I.vH, and E.S. analyzed the data and wrote the manuscript. B.S.V, W.P.L, M.H.I.vH, MM.A.E.C, M.V.A.Q., C.H.I.R, J.E., and E.S. reviewed and edited the manuscript. E.S. supervised the project.

## Declaration of interests

The authors declare no competing interests.

## Notes

### Competing Interest Statement

The authors have declared no competing interest.

