## Supplementary information for "Controlling interfacial protein adsorption, desorption and aggregation in biomolecular condensates"


**This PDF file includes:**

Table of measured zeta potentials and viscosities
Figures S1 to S19
Explanation of fitting of adapted Freundlich model

**Supplementary Table 1: Parameters used to calculate ζ-potentials of pLys/pGlu condensates.** $C_{ion}$ was calculated from the buffer concentration and added sodium chloride. $\eta_d$ was determined by active micro rheology. $E_0$ was determined by testing the lowest electric field strength at which droplets moved. When $\eta_d$ is noted with a *, the viscosity was interpolated or extrapolated. Other parameters used to calculate the ζ-potential are $T$ = 298 K and water viscosity of 0.891 mPa s.

| Sample | $C_{ion}$ (mM) | $\eta_d$ (mPa s) | $E$ (V) | $E_0$ (V) | ζ (mV) |
|---|---|---|---|---|---|
| pLys/pGlu + 0.00 µM αSyn | 127.83 | 80 | 3.0 | 0.5 | +8.1 ± 0.9 |
| pLys/pGlu + 0.10 µM αSyn | 127.83 | 80 | 3.0 | 0.5 | +8.5 ± 0.6 |
| pLys/pGlu + 0.25 µM αSyn | 127.83 | 80 | 3.0 | 0.5 | +5.6 ± 0.9 |
| pLys/pGlu + 0.50 µM αSyn | 127.83 | 80 | 3.0 | 0.5 | +6.3 ± 0.4 |
| pLys/pGlu + 1.00 µM αSyn | 127.83 | 80 | 5.0 | 0.5 | +0.8 ± 0.2 |
| pLys/pGlu + 2.00 µM αSyn | 127.83 | 80 | 6.0 | 0.5 | +3.1 ± 0.4 |
| pLys/pGlu + 3.27 µM αSyn | 127.83 | 80 | 4.0 | 0.5 | +0.9 ± 0.4 |
| pLys/pGlu + 5.00 µM αSyn | 127.83 | 80 | 5.0 | 0.5 | +0.5 ± 0.2 |
| pLys/pGlu + 10.0 µM αSyn | 127.83 | 80 | 6.0 | 0.5 | +0.1 ± 0.0 |
| pLys/pGlu + 20.0 µM αSyn | 127.83 | 80 | 6.0 | 0.5 | +0.1 ± 0.0 |
| pLys/pGlu + 0.5 mM ATP* | 129.33 | 120 | 5.0 | 0.5 | +3.4 ± 0.2 |
| pLys/pGlu + 1.0 mM ATP* | 130.83 | 160 | 5.0 | 1.0 | +2.8 ± 0.2 |
| pLys/pGlu + 3.0 mM ATP | 136.83 | 320 | 3.0 | 1.1 | -1.3 ± 0.5 |
| pLys/pGlu + 5.0 mM ATP* | 142.83 | 480 | 5.0 | 1.3 | -3.0 ± 0.7 |
| pLys/pGlu + 10 mM ATP* | 157.83 | 960 | 5.0 | 1.3 | -2.8 ± 0.4 |
| pLys/pGlu 50 mM NaCl | 77.83 | 100 | 3.0 | 1.0 | +7.5 ± 0.4 |
| pLys/pGlu 200 mM NaCl | 227.83 | 80 | 3.0 | 0.8 | +5.1 ± 0.6 |
| pLys/pGlu 250 mM NaCl | 277.83 | 70 | 3.0 | 0.8 | +6.3 ± 0.4 |
| pLys/pGlu 1.6:1.0* | 127.83 | 80 | 3.0 | 0.8 | +8.1 ± 0.5 |
| pLys/pGlu 1.4:1.0* | 127.83 | 80 | 3.0 | 0.5 | +8.3 ± 0.5 |
| pLys/pGlu 1.2:1.0* | 127.83 | 80 | 3.0 | 0.5 | +8.0 ± 0.4 |
| pLys/pGlu 1.0/1.2* | 127.83 | 80 | 3.0 | 0.5 | +2.4 ± 0.2 |
| pLys/pGlu 1.0/1.4* | 127.83 | 80 | 3.0 | 0.5 | +1.3 ± 0.1 |
| pLys/pGlu 1.0/1.6* | 127.83 | 80 | 3.0 | 0.5 | +0.0 ± 0.0 |
| pLys/pGlu 1.0/2.0* | 127.83 | 80 | 6.0 | 0.5 | -1.3 ± 0.3 |
| pLys/pGlu + 0.1 mM $A_{15}$* | 129.33 | 480 | 5.0 | 1.0 | -4.0 ± 1.0 |
| pLys/pGlu + 0.1 mM $U_{15}$* | 129.33 | 480 | 7.0 | 1.0 | -5.4 ± 0.6 |
| pLys/pGlu + 0.1 mM $C_{15}$* | 129.33 | 480 | 7.0 | 1.0 | -5.4 ± 0.4 |
| pLys/pGlu + 5.0 mM GTP* | 142.83 | 480 | 7.0 | 1.0 | -3.5 ± 0.2 |
| pLys/pGlu + 5.0 mM UTP* | 142.83 | 480 | 10.0 | 1.0 | +0.5 ± 0.4 |
| pLys/pGlu + 5.0 mM CTP* | 142.83 | 480 | 7.0 | 1.0 | -1.3 ± 0.3 |
| pLys/pGlu + 2.5 µg $mL^{-1}$ 1:1* | 127.83 | 80 | 4.0 | 1.0 | +6.5 ± 0.6 |
| pLys/pGlu + 6.3 µg $mL^{-1}$ 1:1* | 127.83 | 80 | 5.0 | 1.0 | +4.7 ± 0.6 |
| pLys/pGlu + 10 µg $mL^{-1}$ 1:1* | 127.83 | 80 | 8.0 | 1.0 | +2.7 ± 0.2 |
| pLys/pGlu + 2.5 µg $mL^{-1}$ 1:2:1* | 127.83 | 80 | 5.0 | 1.0 | +5.6 ± 0.4 |
| pLys/pGlu + 6.3 µg $mL^{-1}$ 1:2:1* | 127.83 | 80 | 5.0 | 1.0 | +3.8 ± 0.3 |
| pLys/pGlu + 10 µg $mL^{-1}$ 1:2:1* | 127.83 | 80 | 5.0 | 1.0 | +1.2 ± 0.4 |
| pLys/pGlu + malate* | 137.83 | 80 | 5.0 | 0.5 | +4.4 ± 0.3 |
| pLys/pGlu + succinate* | 137.83 | 80 | 5.0 | 0.5 | +5.3 ± 0.4 |
| pLys/pGlu + citric acid* | 143.83 | 80 | 7.0 | 0.5 | +4.8 ± 0.2 |
| pLys/pGlu + NADH* | 137.83 | 80 | 5.0 | 0.5 | +3.3 ± 0.3 |
| pLys/pGlu + PPi* | 147.83 | 80 | 9.0 | 0.5 | +0.1 ± 0.0 |
| pLys/pGlu + 0.25 µM eGFP-NPM1 | 127.83 | 80 | 12 | 1.3 | +1.6 ± 0.4 |
| pLys/pGlu + 0.50 µM eGFP-NPM1 | 127.83 | 80 | 15 | 1.3 | +0.1 ± 0.2 |
| pLys/pGlu + 2.50 µM eGFP-NPM1 | 127.83 | 80 | 4 | 1.3 | -0.6 ± 0.1 |
| pLys/pGlu + 5.00 µM eGFP-NPM1 | 127.83 | 80 | 4 | 1.3 | +0.7 ± 0.2 |

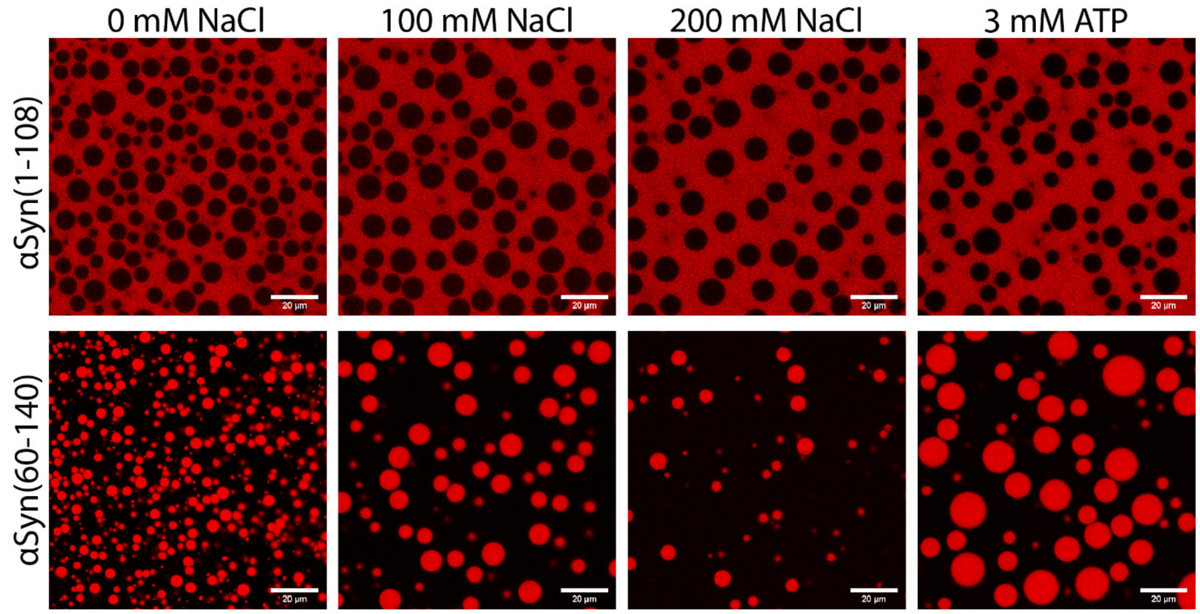


**Supplementary Figure 1: Microscopy of pLys/pGlu condensates with αSyn(1-108) or αSyn(60-140) and various NaCl concentrations and 3 mM ATP.** No substantial changes in partitioning are observed. (Scale bar = 20 µm)

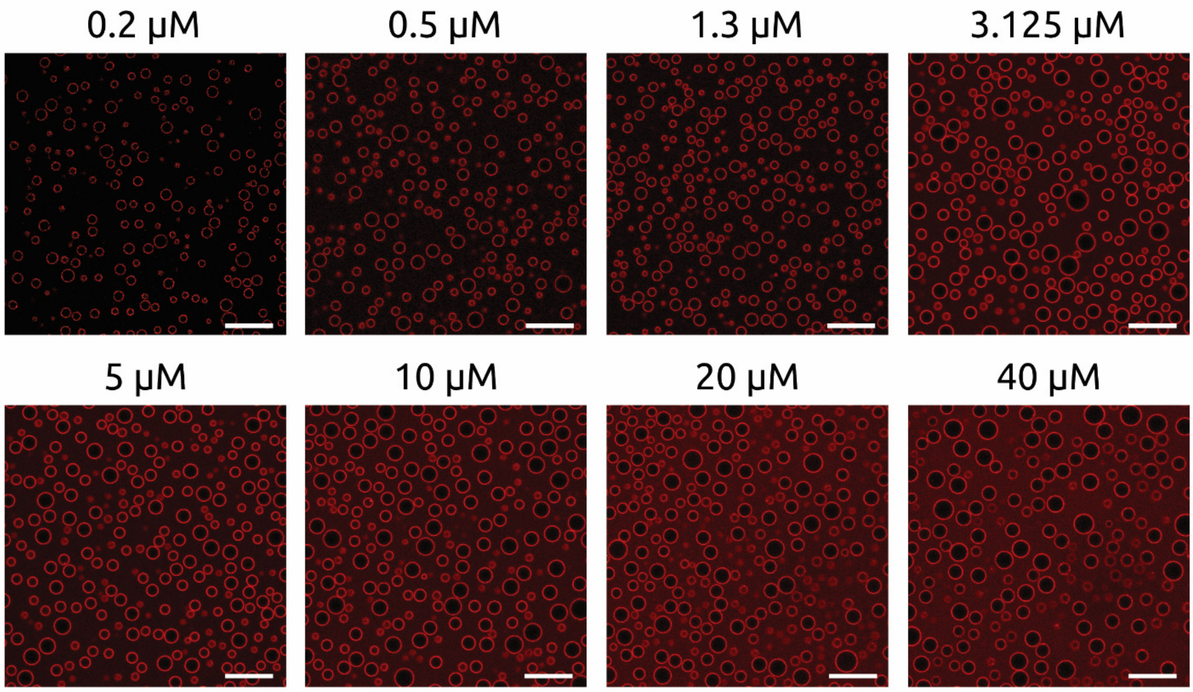


**Supplementary Figure 2: Microscopy of pLys/pGlu condensates with increasing concentrations of αSyn.** (Scale bar = 20 µm)

**Supplementary note 1: Fitting of the adapted Freundlich model to the data obtained from fluorescence microscopy**

From the confocal microscopy images, we can determine the ratio of fluorescence signal at the interface (maximum of the intensity profile along the condensate radius) to the fluorescence signal in the dilute phase. We call this ratio the interface partition coefficient ($K_i$). Technically $K_i$ depends on the ratio of concentration of labelled protein in the confocal volume which includes part of the condensate interface and concentration of labelled protein in the confocal volume in the dilute phase, and it is related to the density of protein molecules at the interface. We used adapted Freundlich isotherm to describe $K_i$ in the function of total protein concentration:

$$K_i = 1 + \frac{\left(\frac{[S]_{\mathrm{dil}}}{[S]_{\mathrm{sat}}}\right)^{1/n}}{\frac{[S]_{\mathrm{dil}}}{[S]_{\mathrm{sat}}}}$$

where: $[S]_{\mathrm{dil}}$ is the concentration in the dilute phase (we assume that the concentration in the dilute phase is equal to the total concentration, as only small fraction of the total protein amount is bound to the interface); $[S]_{\mathrm{sat}}$ is the cross-over concentration, at which the affinity for the interface equals the affinity for the dilute solution, which gives an indication of the adsorption capacity; $n$ is the intensity factor, which gives an indication of the adsorption strength.

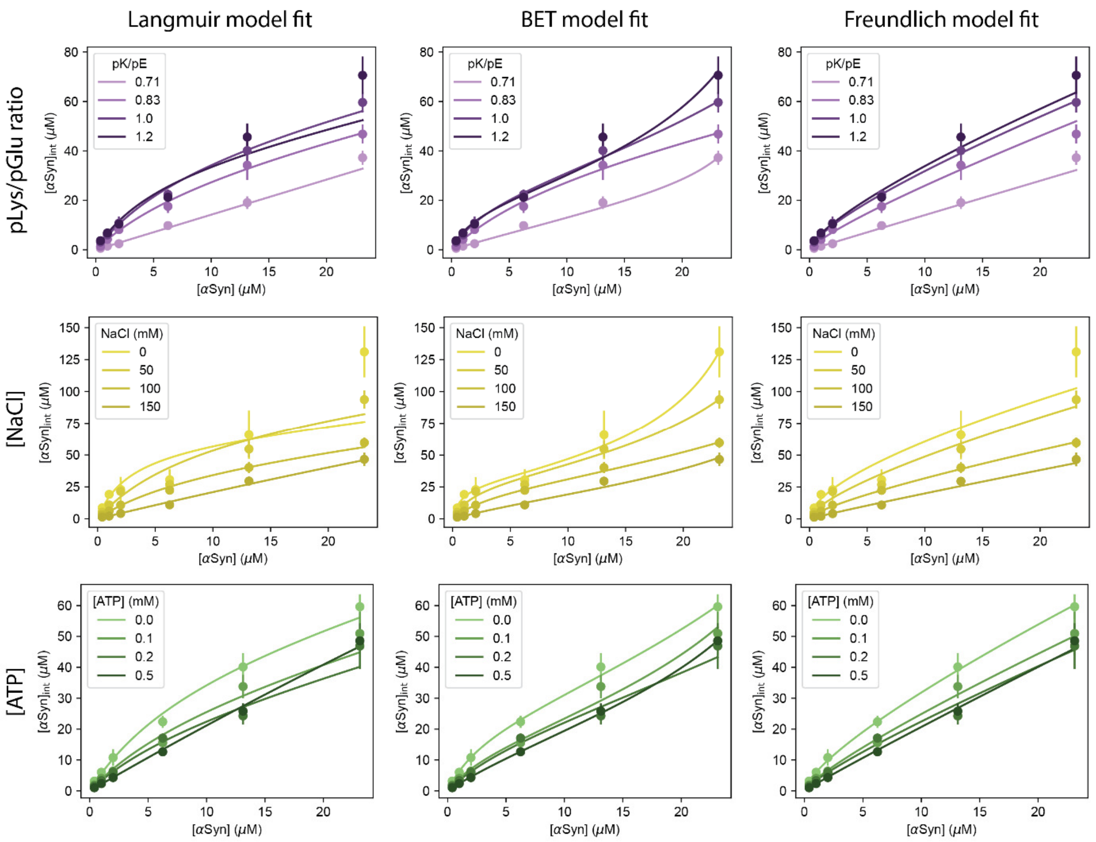


**Supplementary Figure 3: Comparison between adsorption models.** Langmuir, BET and Freundlich model fits for αSyn adsorption for different conditions studied (varying pLys/pGlu ratio, NaCl, and ATP concentration).

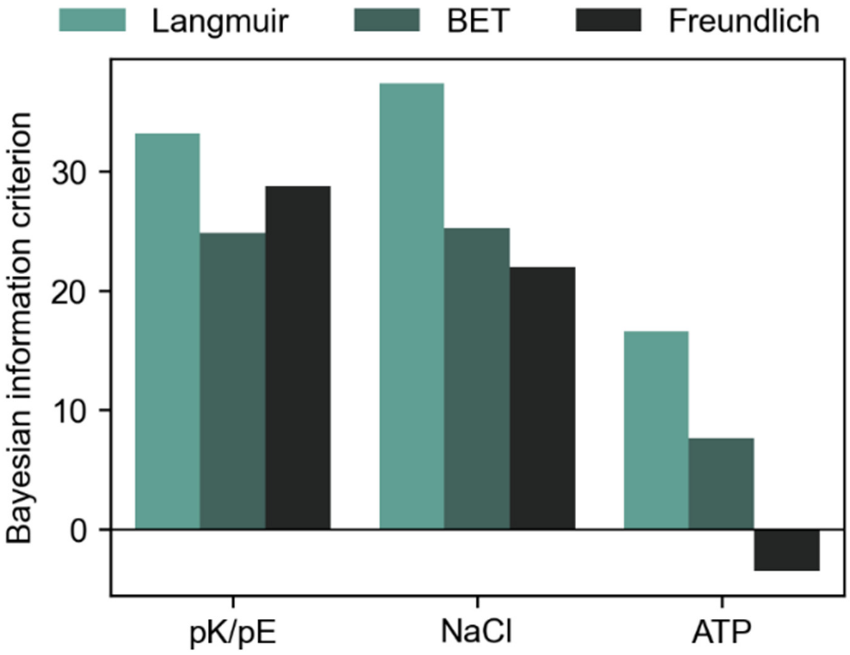


**Supplementary Figure 4: Bayesian Information Criterion (BIC) for the different models in the varying conditions.** Lower BIC values indicate a more suitable model.

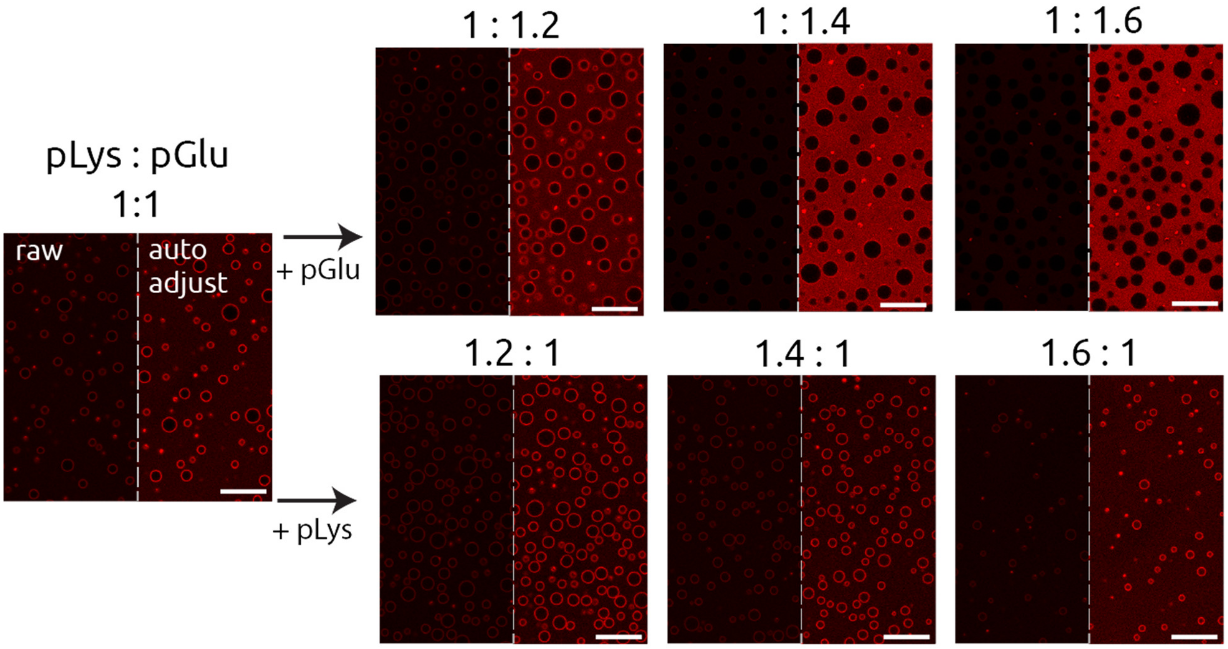


**Supplementary Figure 5: Microscopy of various ratios of pLys/pGlu condensates with αSyn.** FL-αSyn accumulates on the interface of the condensates for ratios smaller than 1:1.4. Raw images are shown on the left, auto-adjusted brightness images on the right. Scale bar = 20 µm.

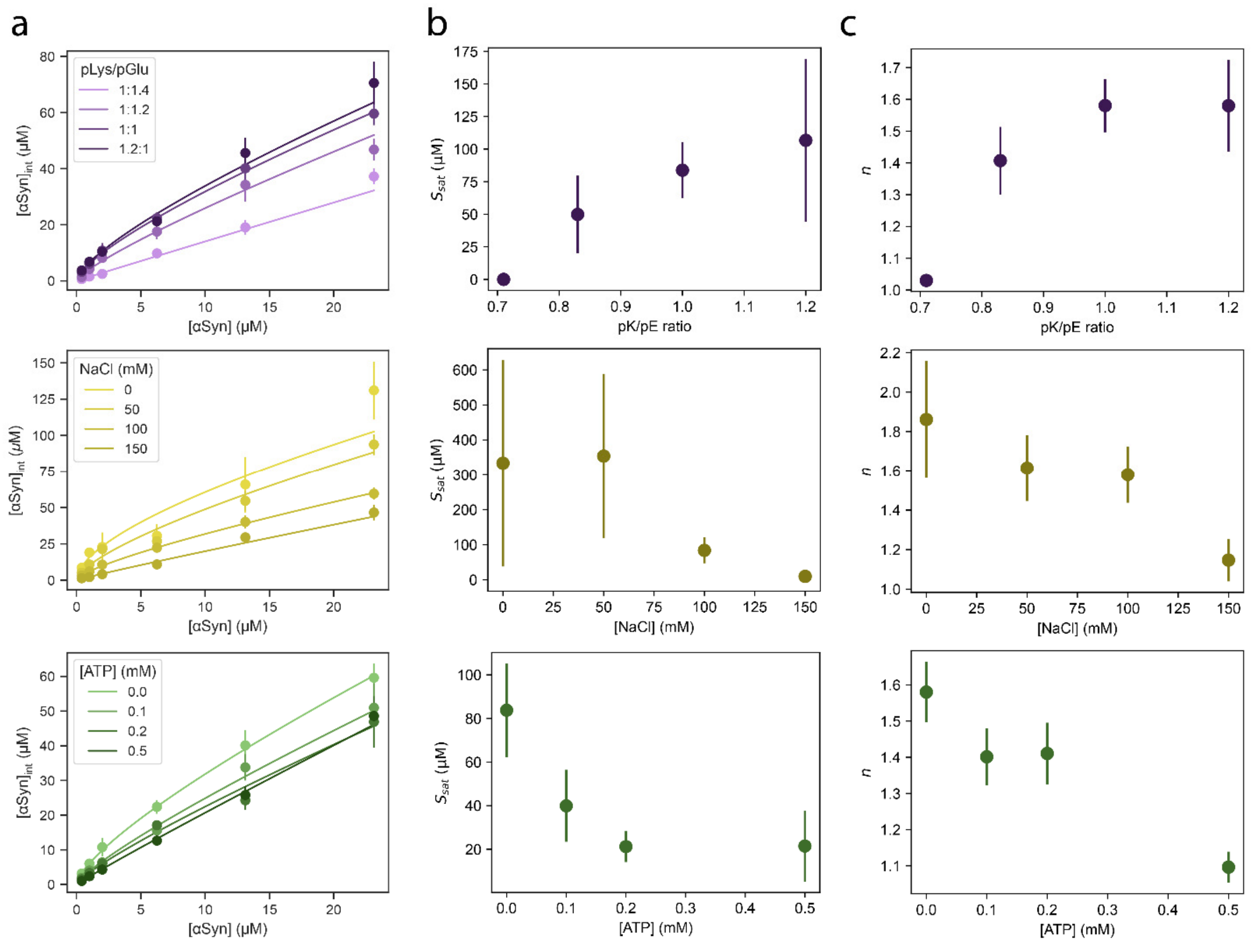


**Supplementary Figure 6: Comparison of Freundlich model parameters for different conditions.** **(a)** Adapted Freundlich model fit to experimentally determined interfacial αSyn concentration for a range of total αSyn concentrations, plotted for different modifier concentrations/component ratios. **(b)** $S_{sat}$ parameter determined from the fit vs. the modifier concentrations/component ratio. **(c)** n parameter determined from the fit vs. the modifier concentrations/component ratio. Upper row: changing pLys/pGlu ratio; middle row: NaCl as the modifier; lower row: ATP as the modifier.

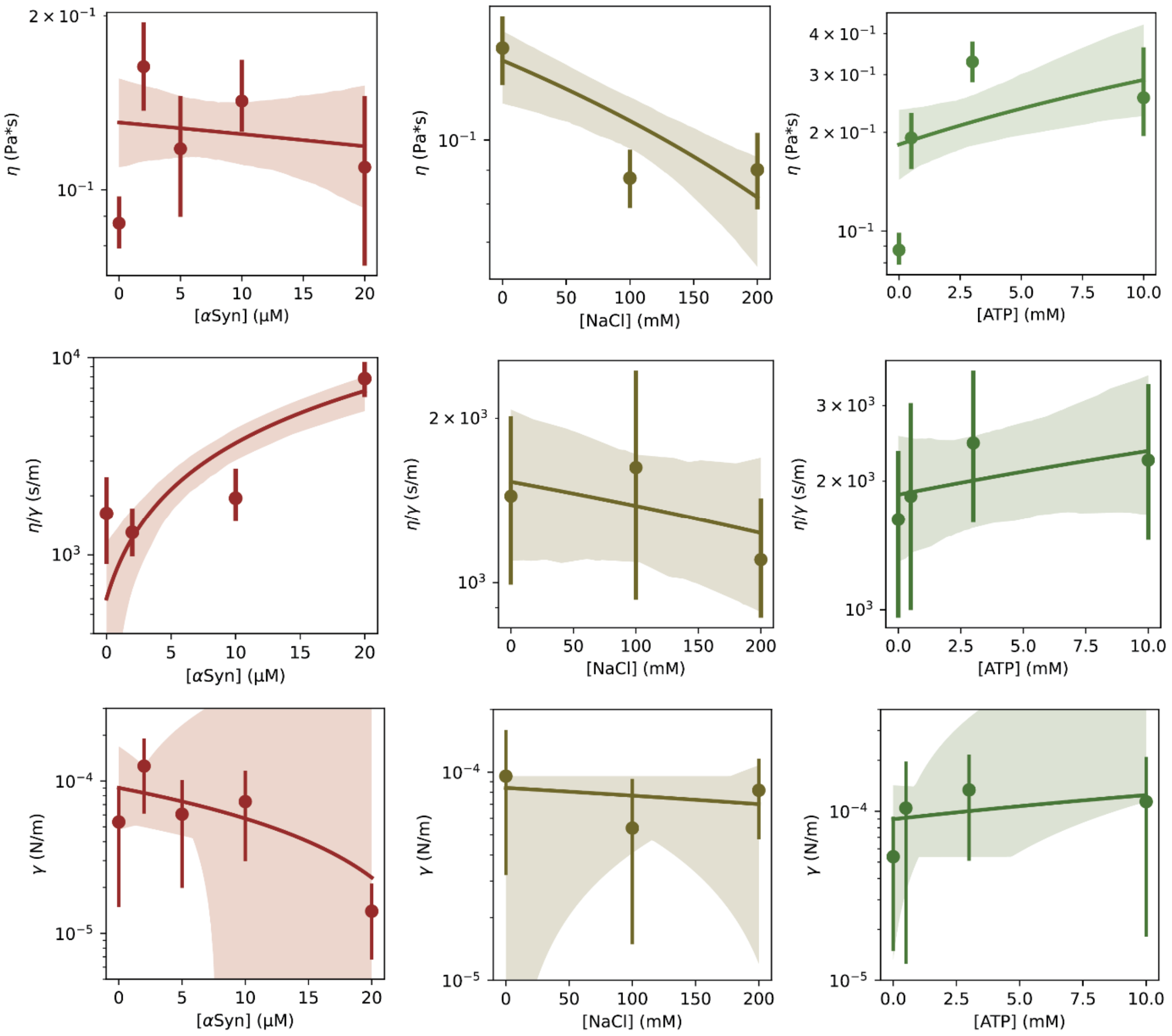


**Supplementary Figure 7: Optical tweezers measurements of physical properties of condensates.** Measured viscosity, viscosity/surface tension ratio, and calculated surface tension of pLys/pGlu upon addition of αSyn, ATP, and NaCl.

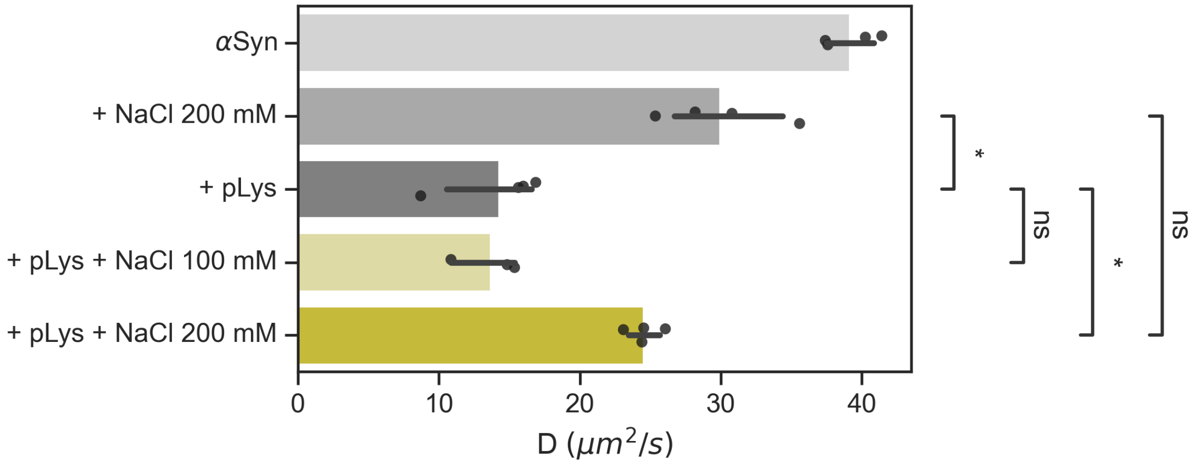


**Supplementary Figure 8: Diffusion coefficients of αSyn in the presence of pLys and NaCl.** The diffusion coefficient of αSyn decreases significantly with the addition of pLys. The introduction of 200 mM NaCl, but not 100 mM NaCl, restores the diffusion coefficient of αSyn to levels observed in the absence of pLys.

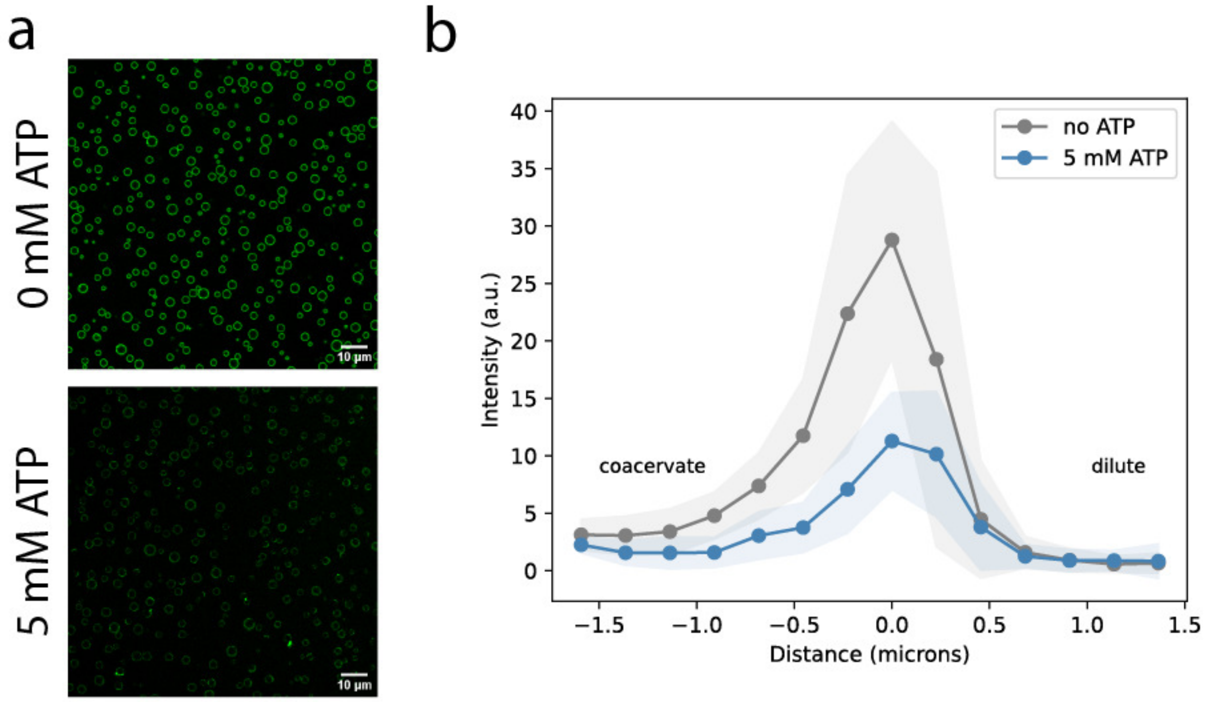


**Supplementary Figure 9: TDP-43-TEV-mCherry can also be removed from the interface of pLys/pGlu condensates by addition of ATP. (a)** TDP-43-TEV-mCherry localizes to the interface of pLys/pGlu condensates. Addition of ATP partially removes it from the interface. Microscopy images are shown at the same brightness/contrast for comparison. **(b)** Radial intensity plot of TDP-43 at the interface of pLys/pGlu show that addition of ATP reduces the interfacial localization.

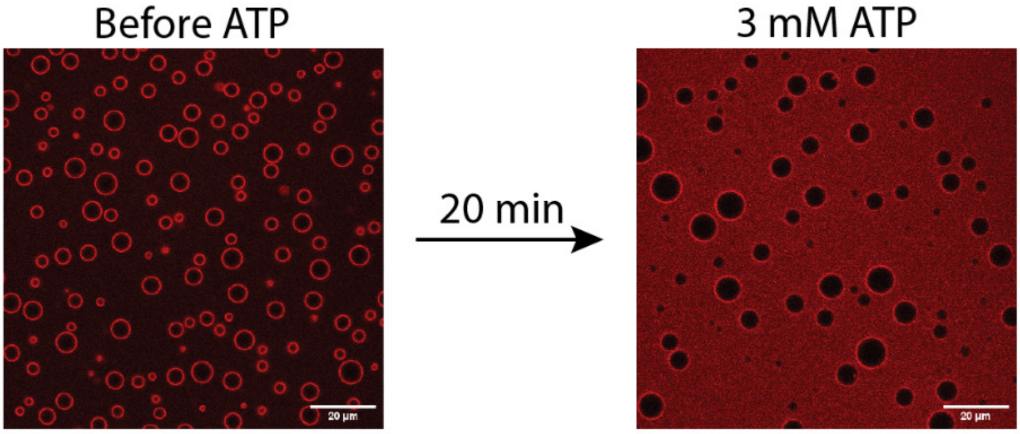


**Supplementary Figure 10: Addition of ATP to pLys/pGlu condensates with interfacially localized αSyn can remove αSyn from the interface.** (Scale bar = 20 µm)

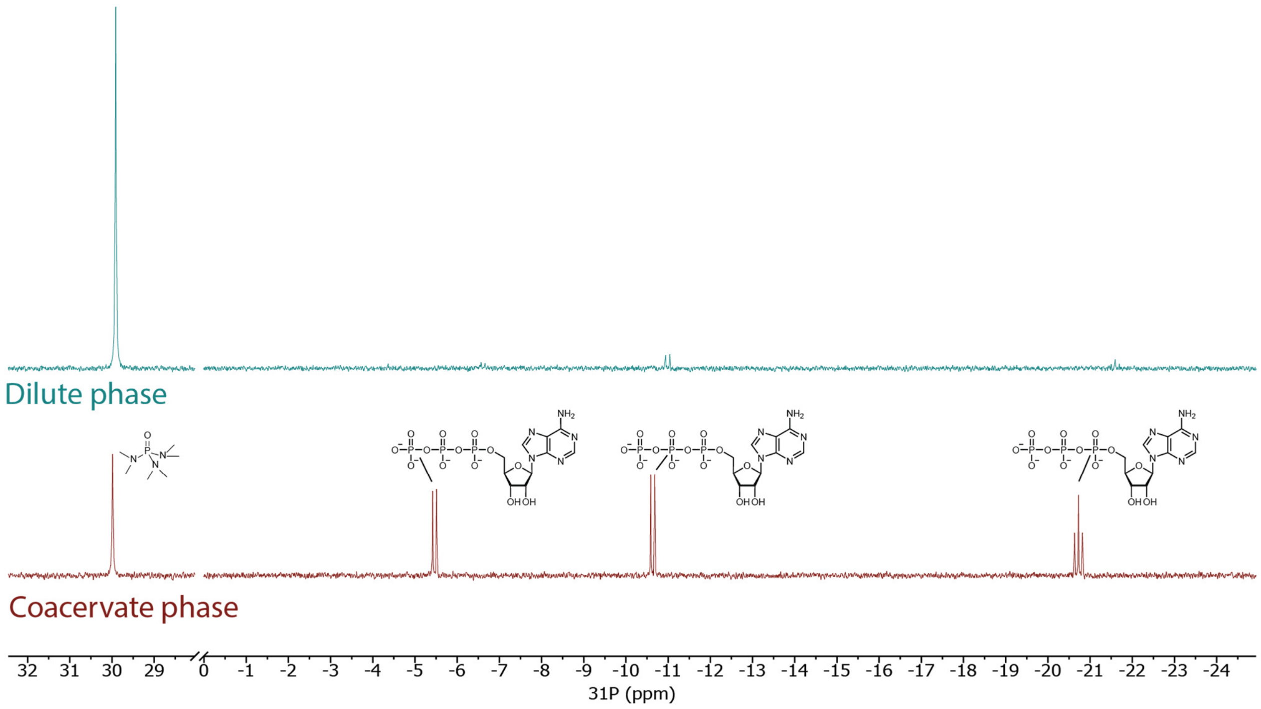


**Supplementary Figure 11**: $^{31}$P-NMR spectra of ATP in the separated condensate phase (bottom, red) and dilute phase (top, blue). Dilute phase contains 10 mM HMPA internal standard, whereas condensate phase contains 4 mM HMPA.

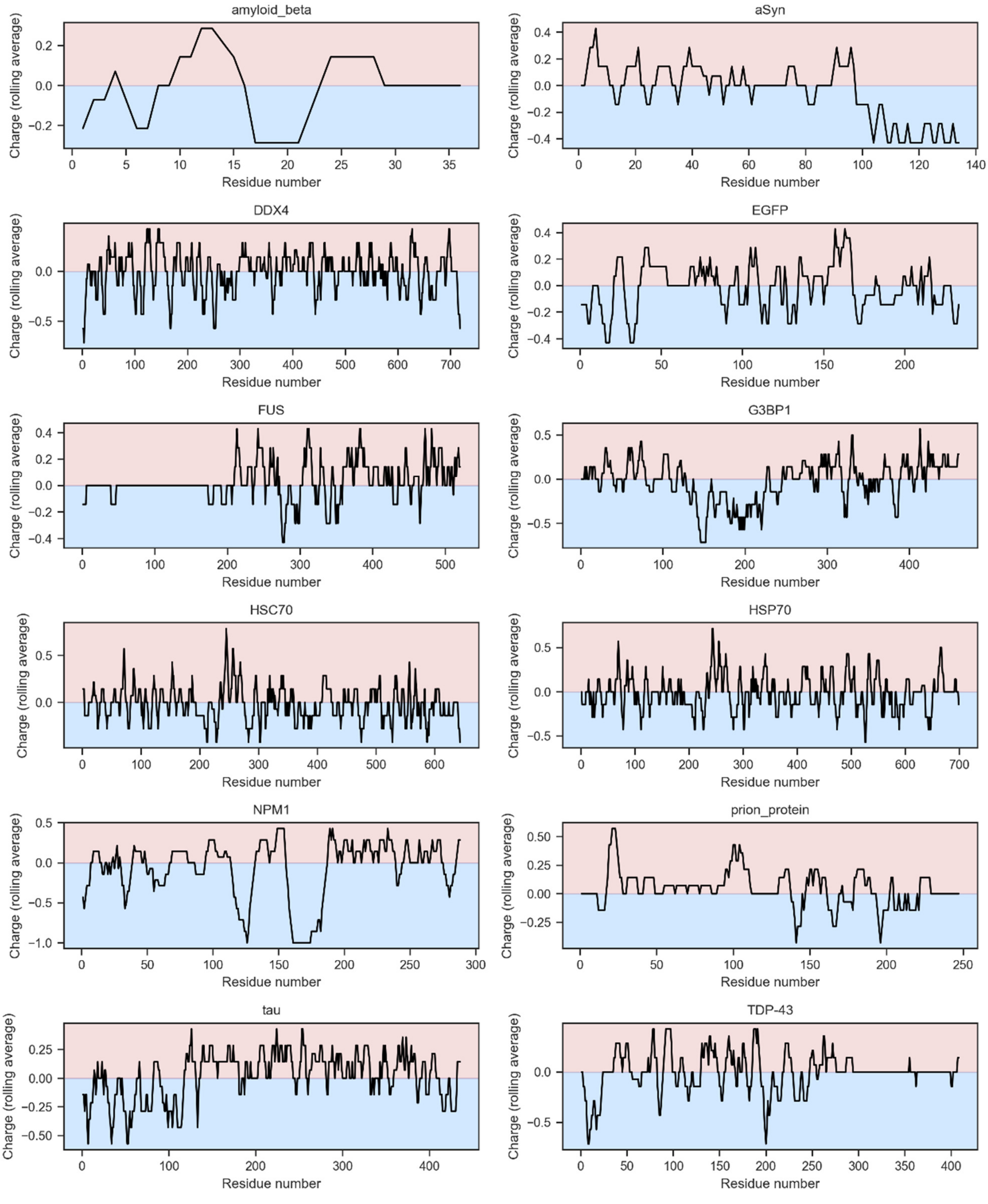


**Supplementary Figure 12:** Charged patches (rolling average with window 11) are present in various aggregation-related proteins, and in other proteins used in this study.

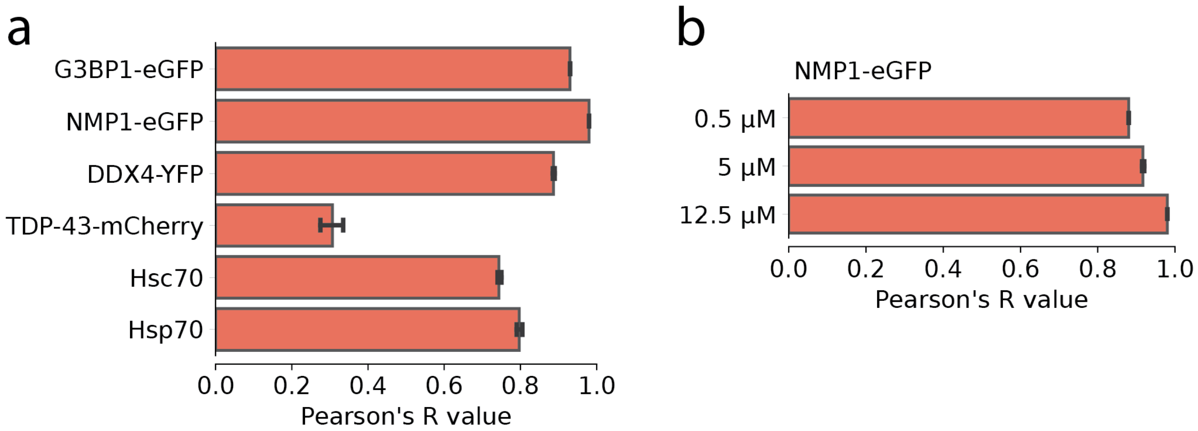


**Supplementary Figure 13:** **(a)** Colocalization of added protein and αSyn on the interface of pLys/pGlu condensates. **(b)** Colocalization of NPM1-EGFP increases with increasing concentrations of NPM1.

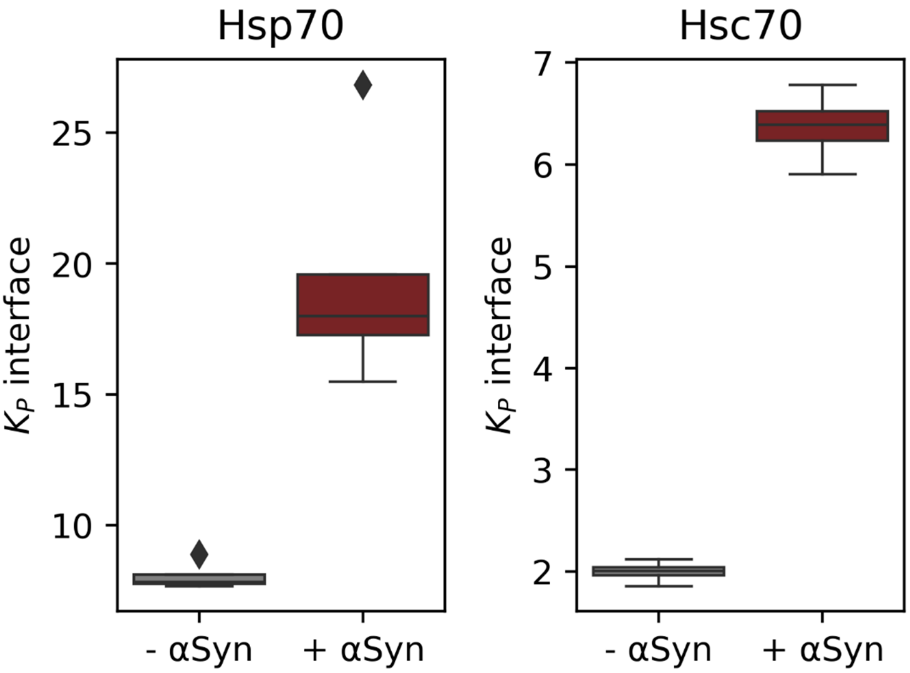


**Supplementary Figure 14:** chaperones localize to the pLys/pGlu interface more strongly when αSyn is present.

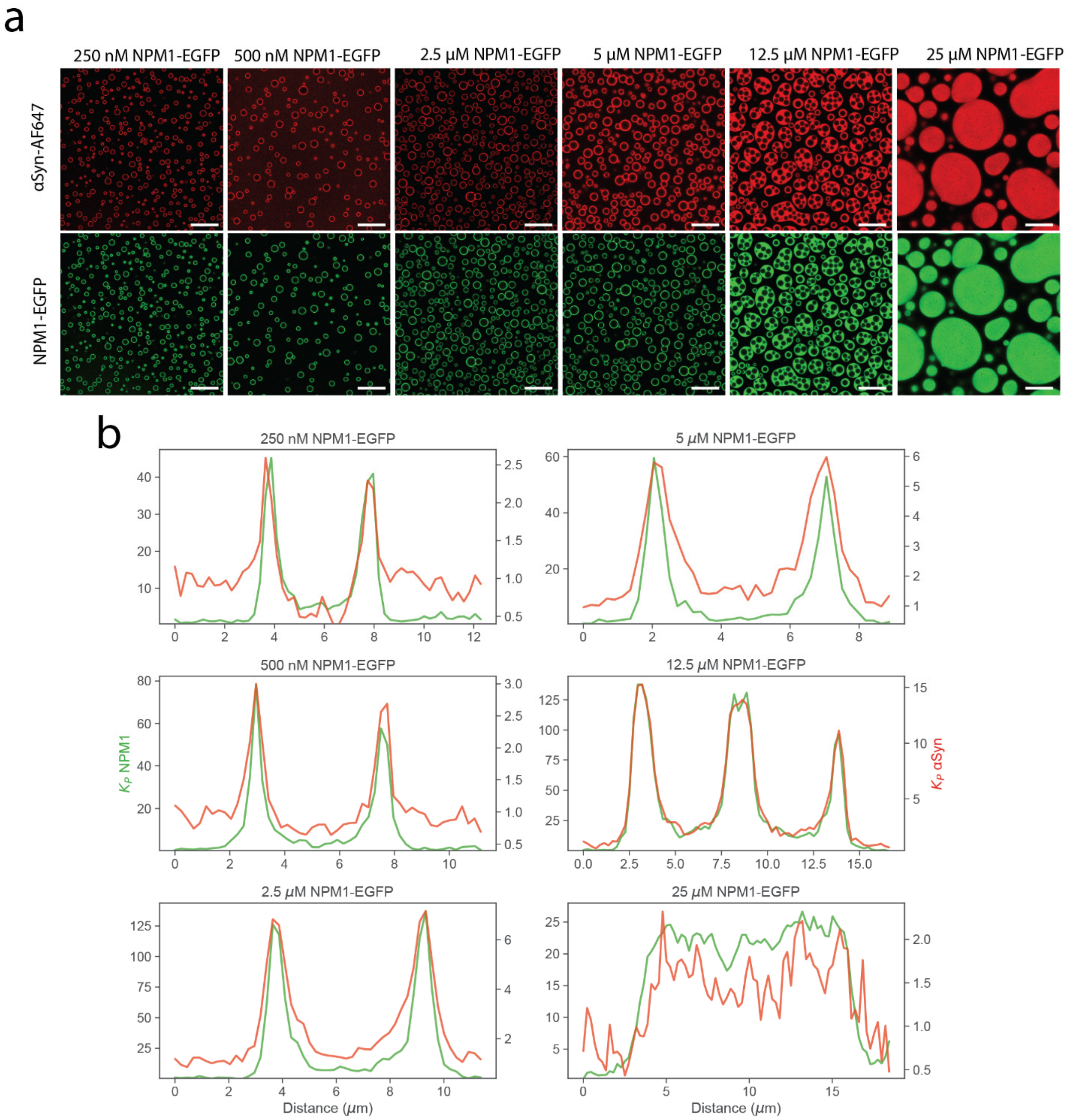


**Supplementary Figure 15: αSyn and NPM1-eGFP localization with pLys/pGlu condensates. (a)** Microscopy of various concentrations of NPM1-EGFP added to pLys/pGlu condensates with αSyn. NPM1 partitions to the interface and at 12.5 μM forms multiphasic condensates. (Scale bar = 20 μm) **(b)** Normalized fluorescence intensity profiles. NPM1 partitioning to the interface increases and phase separates at 12.5 μM. The multiphase condensates become a single phase at higher concentration. αSyn also partitions higher with increasing NPM1, with the highest partitioning in the multiphase condensates. Intensity is normalized to dilute phase signal and a single line was taken for the profile.

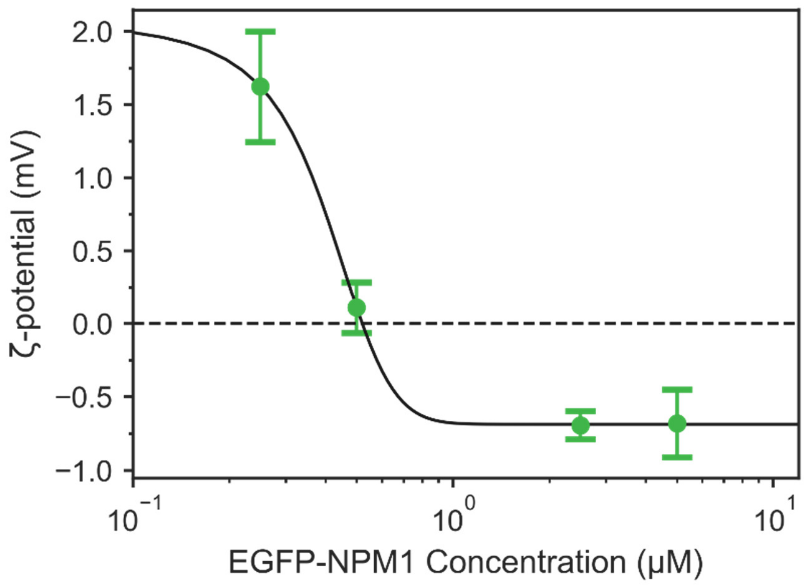


**Supplementary Figure 15:** ζ-potential of pLys/pGlu condensates with increasing concentrations EGFP-NPM1**.**

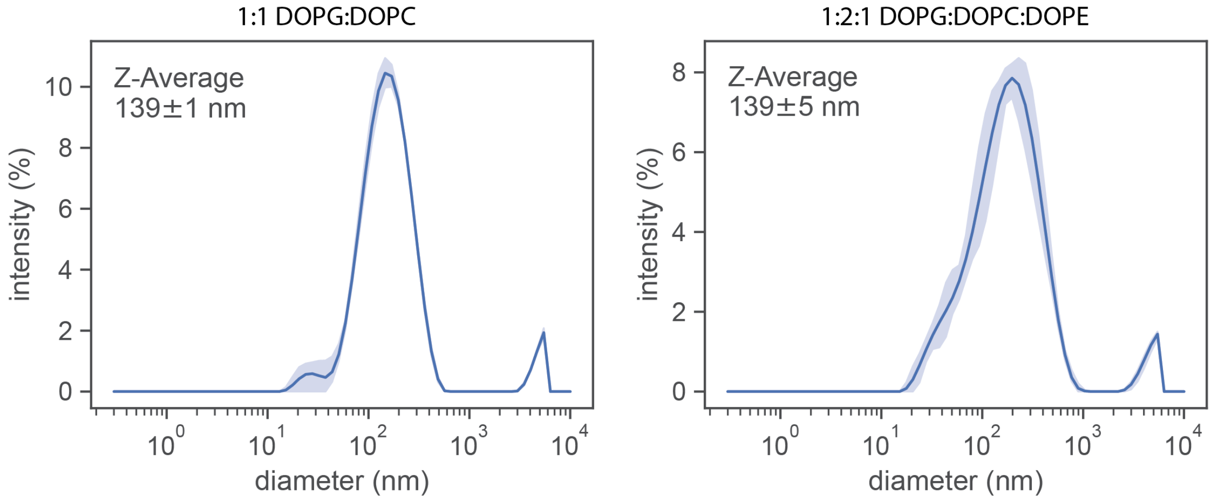


**Supplementary Figure 16: DLS determined size distribution (intensity based) of lipid vesicles used in the study.**

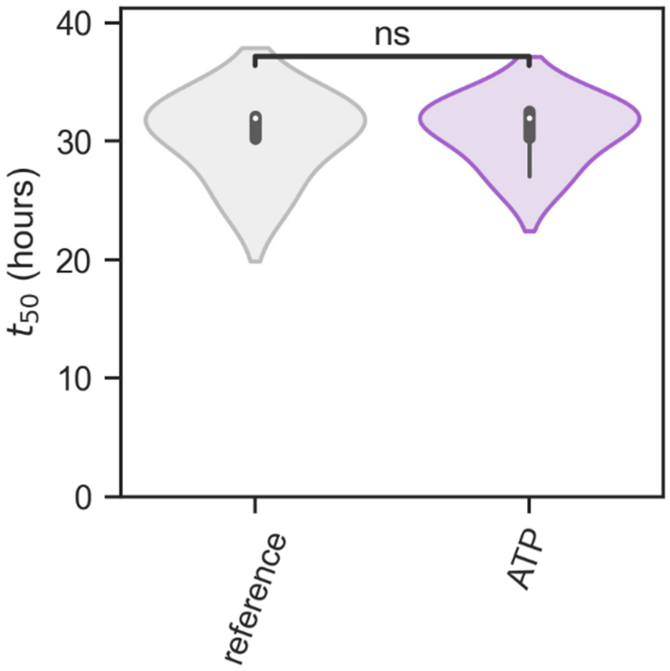


**Supplementary Figure 17: αSyn aggregation with and without 5 mM ATP in the absence of condensates.** No significant difference is observed (Welch's t-test independent samples, $p = 0.84$, n=4).

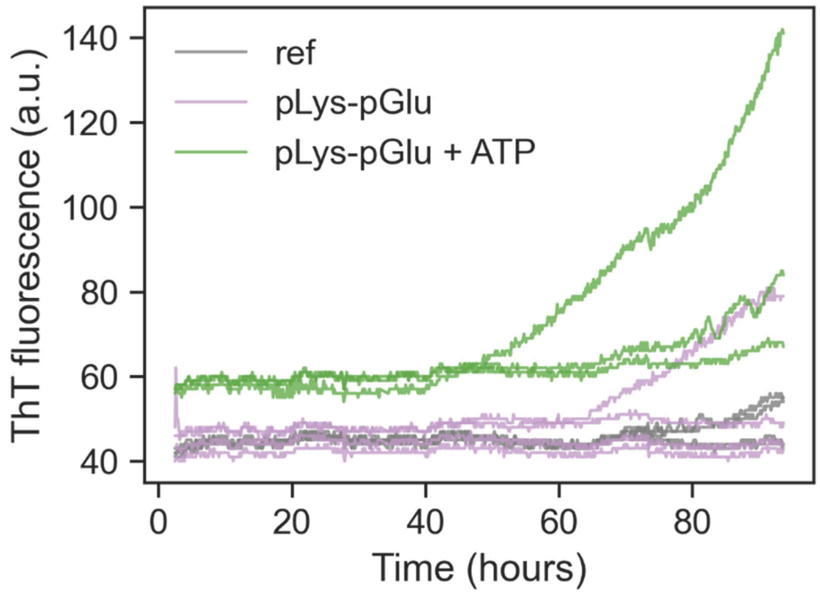


**Supplementary Figure 18: ThT fluorescence of αSyn(60-140) (gray, ref) in the presence of pLys-pGlu condensates (purple) and condensates with ATP (green).** No substantial difference in kinetics can be seen, except for a higher baseline ThT fluorescence. Samples were not fully aggregated in the experiment duration, as measured ThT fluorescence is more that 1-2 orders of magnitude smaller than for aggregated samples.

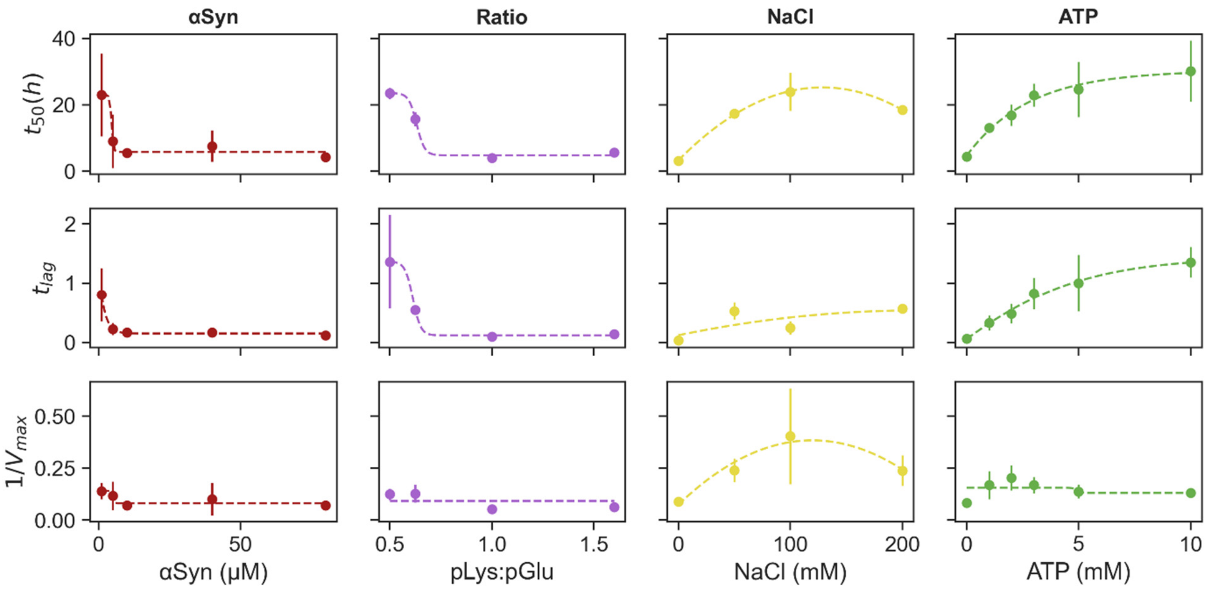


**Supplementary Figure 19: Preventing interfacial localization of αSyn, regardless of method used, slows down aggregation.** Increasing the concentration of αSyn leads to faster aggregation at low concentrations but plateaus due to limited interface availability. Lag times are increased in all methods of interfacial αSyn prevention, while max elongation rates are only reduced when NaCl or ATP is added. Lines are drawn to guide the eye and were generated by fitting either a sigmoidal curve or 2$^{nd}$ order polynomial.
